# Neuroendocrine and behavioral measures of stress-reactivity in male goal-tracker and sign-tracker rats

**DOI:** 10.1101/2020.09.04.283549

**Authors:** Sofia A. Lopez, Eman Mubarak, Charlotte Yang, Aram Parsegian, Marin Klumpner, Paolo Campus, Shelly B. Flagel

## Abstract

Environmental cues attain the ability to guide behavior via learned associations. As predictors, cues can elicit adaptive behavior and lead to valuable resources (*e.g*., food). For some individuals, however, cues are transformed into incentive stimuli and elicit motivational states that can be maladaptive. The goal-tracker/sign-tracker animal model captures individual differences in cue-motivated behaviors, with reward-associated cues serving as predictors of reward for both phenotypes but becoming incentive stimuli to a greater degree for sign-trackers. While these distinct phenotypes are characterized based on Pavlovian conditioned approach behavior, they exhibit differences on a number of behaviors relevant to psychopathology. To further characterize the neurobehavioral endophenotype associated with individual differences in cue-reward learning, neuroendocrine and behavioral profiles associated with stress and anxiety were investigated in male goal-tracker, sign-tracker, and intermediate responder rats. It was revealed that baseline corticosterone increases with Pavlovian learning, but to the same degree, regardless of phenotype. No significant differences in behavior were observed between goal-trackers and sign-trackers during an elevated plus maze or open field test, nor were there differences in corticosterone response to the open field test or physiological restraint. Upon examination of central markers associated with stress-reactivity, we found that sign-trackers have greater glucocorticoid receptor mRNA expression in the ventral hippocampus, with no phenotypic differences in the dorsal hippocampus or prelimbic cortex. These findings demonstrate that goal-trackers and sign-trackers do not differ on stress- and anxiety-related behaviors, and suggest that differences in neuroendocrine measures between these phenotypes can be attributed to distinct cue-reward learning styles.

**Significance Statement:** While the goal-tracker/ sign-tracker animal model derives from individual differences in Pavlovian conditioned approach behavior, other traits, including some of relevance to addiction and post-traumatic stress disorder, have been shown to co-exist with the propensity to sign-track. The extent to which this model encompasses differences in aversive arousal and associated neuroendocrine measures, however, remains largely unexplored. Here we show that behavioral and corticosterone response to stress-related paradigms do not differ between goal-trackers and sign-trackers. However, glucocorticoid receptor expression in the ventral hippocampus does differ between phenotypes, suggesting that this central marker that is typically associated with stress-responsivity, may, in fact, play an important role in appetitive motivation.

## Introduction

Through learned associations, environmental cues become predictors of biologically relevant stimuli. In turn, such cues elicit an adaptive response, facilitating behavior towards valuable resources. For some individuals, however, cues elicit complex emotional responses and can prompt maladaptive behavior. For example, upon exposure to drug-associated cues, individuals with addiction report drug-craving and, consequently, often relapse (Ehrman et al., 1992). Similarly, when exposed to trauma-related stimuli, individuals with post-traumatic stress disorder (PTSD) report hyperarousal and anxiety (Shin et al., 2004). Cues attain the ability to elicit extreme emotional states and aberrant behavior when they are attributed with excessive incentive motivational value, or incentive salience (Robinson & Berridge, 1993). The propensity to attribute incentive salience to environmental cues, thereby, may reflect a vulnerability trait for cue-motivated psychopathologies, like addiction and PTSD (Flagel et al., 2010; Morrow et al., 2011).

Individual variation in the propensity to attribute incentive salience to reward cues can be captured using a Pavlovian conditioned approach (PavCA) paradigm, consisting of a lever-cue paired with delivery of a food-reward (Robinson & Flagel, 2009). Upon lever-cue presentation, goal-trackers (GTs) direct their behavior towards the location of reward delivery, whereas sign-trackers (STs) approach the cue itself. For both GTs and STs the cue attains predictive value, but for STs the cue also attains incentive value and is transformed into a “motivational magnet” (Berridge et al., 2009). Intermediate responders (IRs) vacillate between goal- and cue-directed behavior, without preference for either cue-learning strategy. GTs and STs differ on a number of traits of relevance to psychopathology. Relative to GTs, STs are more impulsive (Lovic et al., 2011), exhibit an exaggerated fear response to aversive stimuli (Morrow et al., 2011; Morrow et al., 2015), show poor attentional control (Paolone et al., 2013), and have a greater propensity for reinstatement of drug-seeking behavior (Flagel et al., 2010; Saunders & Robinson, 2010, 2011; Saunders et al., 2013; Yager & Robinson, 2013, *also see* Kawa et al., 2016). These behavioral phenotypes are subserved by distinct neural mechanisms (Campus et al., 2019; Flagel, Clark, et al., 2011; Pitchers et al., 2017). While GTs seem to rely on “top-down” cortical control, STs are presumed to be driven by subcortical “bottom-up” circuitry (Flagel & Robinson, 2017; Kuhn et al., 2018; Sarter & Phillips, 2018). Thus, the GT/ST model captures a neurobehavioral endophenotype reflective of more than individual differences in cue-reward learning.

Most of the research surrounding the GT/ST model has focused on appetitive motivation, with only a few studies investigating indices of aversive arousal (*e.g*., Harb & Almeida, 2014; Morrow et al., 2011; Vanhille et al., 2015). Corticosterone (CORT), the final product of the hypothalamic-pituitary-adrenal (HPA) axis in rodents, is recognized as a biomarker of stress (*e.g*., Dallman & Jones, 1973). However, we know that the role of CORT extends into arenas of learning and memory (*e.g*., Sandi et al., 1997), reward-learning (*e.g*., Tomie et al., 2002), and reinforcement (*e.g*., Piazza et al., 1993). Of particular relevance, CORT is involved in forming Pavlovian associations for both aversive (*e.g*., Marchand et al., 2007) and appetitive (*e.g*., Tomie et al., 2002) stimuli (*for review, see* Lopez & Flagel, 2020). With respect to the latter, relative to GTs, STs show a greater rise in CORT following an initial PavCA session, prior to the development of a conditioned response (Flagel et al., 2009). Baseline CORT levels prior to training do not differ between phenotypes (Flagel et al., 2009), but it remains to be determined whether baseline CORT changes as a consequence of cue-reward learning. This was particularly important to assess in the current study, in order to account for potential learning-induced differences in CORT that may alter stress-responsivity in subsequent tests. Thus, in Experiment 1A, we assessed baseline CORT levels before and after the acquisition of PavCA behavior. We hypothesized that any changes in CORT over the course of Pavlovian learning would be a function of the propensity to attribute incentive salience to reward cues. In Experiment 1B, we assessed CORT and behavioral responses to the elevated plus maze, open field test, and acute physiological restraint, to determine whether goal-trackers and sign-trackers differ in stress-responsivity. We hypothesized that these phenotypes would not differ in stress-reactivity and that any differences in CORT would be specific to Pavlovian learning. Further, in Experiment 2, to examine a central regulator of stress-responsivity (Akil, 2005; Reul et al., 1987), we assessed glucocorticoid receptor (GR) mRNA in the hippocampus and prelimbic cortex. These brain regions were selected as both are integral for regulation of the stress response (*for review, see* Herman et al., 2003), and both have been implicated in incentive salience attribution (*e.g*., Campus et al., 2019; Fitzpatrick et al., 2016). Thus, we hypothesized that GR mRNA would differ between phenotypes in the hippocampus and prelimbic cortex. Together, these studies expand the characterization of the neurobehavioral endophenotype captured by the GT/ST model.

## Materials and Methods

### Experiment 1: General procedures

#### Animals

For Experiment 1(A-B), sixty male Sprague-Dawley rats were obtained from Charles River Breeding Labs (Colony 72 (C72), Saint-Constant, Canada, and Colony 04 (R04), Raleigh, NC, USA). Rats weighed between 225-275 g upon arrival and were pair-housed in standard acrylic cages (46 x 24 x 22 cm) in a temperature-controlled room (22 ± 2°C) under a 12-h light/dark cycle (lights on at 7:00). Food and water were available ad libitum for the duration of the study. Rats were allowed to acclimate to their colony room and remained undisturbed in their homecages for seven days after arrival. Rats were then briefly handled every day for five consecutive days before any experimental manipulation. During the last two days of handling, twenty-five 45-mg banana-flavored grain pellets (Bio-Serv, Flemington, NJ, USA) were placed inside the homecage, allowing rats to habituate to the food reward used during Pavlovian conditioned approach (PavCA) training. Behavioral testing occurred during the light cycle (between 10:00 and 14:00). All experimental procedures followed The Guide for the Care and Use of Laboratory Animals: Eight Edition (2011, National Academy of Sciences), and were approved by the University of Michigan Institutional Animal Care and Use Committee.

#### Behavioral testing

##### Pavlovian conditioned approach (PavCA) training

All PavCA training took place in standard behavioral testing chambers (MED Associates, St. Albans, VT, USA; 20.5×24.1 cm floor area, 29.2 cm high) located inside a room with red lighting. The chambers were enclosed in sound-attenuating boxes equipped with a ventilation fan that provided constant air circulation and served as white noise. Each chamber contained a foodcup centered on one of the walls and placed 6 cm above the grid floor. The food-cup was equipped with an infrared beam, and each beam break was recorded as a head entry. Counterbalanced, right or left of the food-cup, was a retractable lever that illuminated upon presentation and was also placed 6 cm above the floor. A force of at least 10 g was necessary to deflect the lever; this deflection was recorded as a “lever contact.” On the opposite wall, a white house light was placed 1 cm from the top of the chamber. House light illumination signaled the beginning of the session and remained on for the duration of the session.

Rats underwent a single pre-training session, where the food-cup was baited with three grain pellets in order to direct the rat’s attention to the location of the reward. Once placed in the chamber, the house light turned on after 5 min, signaling the beginning of the session. The pretraining session consisted of 25 trials during which the lever remained retracted, and pellets were delivered randomly into the food-cup; one pellet per trial on a variable interval 30 s schedule (range 0-60 s). The total session length was approximately 12.5 min.

Following pre-training, or twenty-four hours later, rats underwent a total of five consecutive PavCA training sessions. Each session consisted of 25 trials on a variable interval 90 s schedule (VI 90, range 30-150 s) during which an illuminated lever (conditioned stimulus, CS) was presented for a total of 8 s, and immediately upon its retraction, a food pellet (unconditioned stimulus, US) was delivered into the adjacent food-cup. Each session lasted approximately 40 min.

The following behavioral measures were recorded during each PavCA session: (1) number of lever contacts, (2) latency to contact the lever for the first time, (3) probability to contact the lever, (4) number of food-cup entries during presentation of the lever, (5) latency to first enter the food-cup during presentation of the lever, (6) probability of entering the food-cup during presentation of the lever, and (7) number of food-cup entries during the inter-trial interval. These values were then used to calculate three measures of approach behavior that comprise the PavCA index: (1) response bias [(total lever presses – total food-cup entries) ÷ (total lever presses + total food-cup entries)], (2) probability difference [probability to approach the lever – the probability to enter the food-cup], (3) latency difference [± (latency to approach the lever – latency to enter the food-cup) ÷ 8]. As previously described (Meyer et al., 2012), PavCA index score was calculated from the average of sessions 4 and 5 using this formula: [(response bias + probability difference + latency difference) ÷ 3]. Scores ranged from +1 to −1; a more positive score indicated a preference for sign-tracking behavior and a negative score for goal-tracking. The cutoffs for phenotype classification were: ≤ −0.5 for a GT, ≥ 0.5 for a ST, and in between - 0.5 and 0.5 for an IR, those that vacillate between the two conditioned responses.

#### Corticosterone

##### Sample collection

To investigate plasma corticosterone (CORT) profiles, blood samples were collected via lateral tail nick at the time points indicated below for Experiment 1A and 1B (*see also* Figure 1A). An experimenter lightly restrained each rat under a blue pad near the edge of a flat surface, allowing their tail to hang off. A small (≤ 5 mm) nick was made with the tip of a razor blade, and blood was extracted via capillary action (~200 μL) into an EDTA-coated tube (Sarstedt, Nümbrecht, Germany). Samples were capped, inverted 2-3 times, and immediately placed onto ice where they remained until the last tail nick was performed (< 3 hr standing time). Samples were then separated by centrifugation (13,000 rpm for 10 min at 4 °C), and plasma was extracted, flash-frozen on dry ice, and stored at −20 °C until processed for radioimmunoassay.

**Figure 1.**
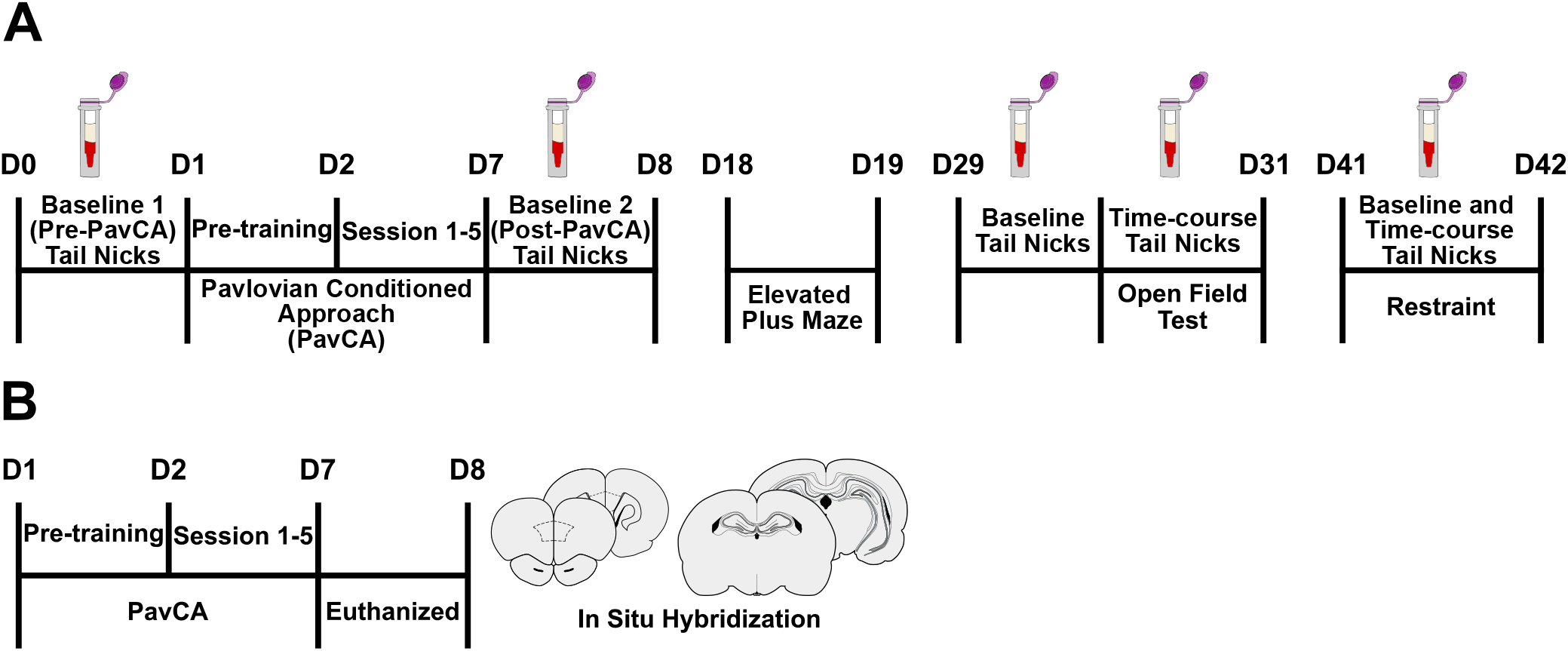
Experimental Timelines. **A**) “Baseline” tail nicks were performed for blood collection prior to Pavlovian conditioned approach (PavCA) training (Pre-PavCA), and after the rats had acquired a conditioned response (Post-PavCA). Rats were subsequently tested on an elevated plus maze (EPM) and the open field test (OFT), followed by physiological restraint, with a 10-day rest period prior to each. Corticosterone response to the OFT and acute restraint was captured with time-course blood sampling. **B**) A separate group of rats underwent 5 sessions of PavCA training and were subsequently euthanized to assess glucocorticoid receptor expression in the hippocampus and prelimbic cortex using in situ hybridization.

##### Radioimmunoassay

Plasma CORT levels were measured using commercially available CORT I^125^ Double Antibody Radioimmunoassay (RIA) kit (MP Biomedicals, Solon, OH) with a minimum detectable dose of 7.7 ng/mL. The manufacturer’s protocol was followed verbatim (*and as reported in* Clinton et al., 2014). A range of 25-1000 ng/mL CORT calibrators was used to generate a standard logarithmic curve for every set of 76 test tubes (the centrifuge test tube capacity for one spin). For Experiments 1A and 1B, a total of 482 plasma samples (not including duplicates or calibration standards) were assayed using 19 centrifuge spins across 6 days, with no more than 4 sets (*i.e*., centrifuge spins) per day. Gamma radiation counts per minute were averaged across duplicate samples and converted into CORT concentrations using the average standard curve generated from all sets that were run for each day of RIA. On average, the intraassay coefficient of variation was 7.24%, while, the inter-assay coefficient of variation was 16.44%. Outliers were identified and removed if: 1) duplicates had a percent error greater than 10%, or 2) samples were identified as an extreme outlier (3x the interquartile range) by statistical software.

### Experiment 1A: Pavlovian conditioned approach behavior and baseline plasma corticosterone profiles

#### Corticosterone

##### Sample collection

Samples were collected, as described above, under baseline conditions before Pavlovian conditioned approach training (Pre-PavCA) and following the development of a conditioned response to the lever-cue (Post-PavCA) (*refer to the experimental timeline*, Figure 1A). Pre-PavCA tail nicks were performed 24 h prior to the pre-training sessions (*see* Experiment 1 Behavioral Testing), while Post-PavCA tail nicks were performed 24 h after the last session (Session 5) of training. On days of collection, six rats were transported in their paired-housed homecages into a designated room (start 10:30), where all collection took place under white light. Tail nicks were performed one at a time (~ 60-90 s per collection). Each wave of six rats remained in the room together but on the opposite side of the room from the collection area. After the last tail nick was performed, all rats were promptly returned to the colony room. Rats were left undisturbed for a total of ten-days before Experiment 1B began.

### Experiment 1B: Behavioral and corticosterone response to anxiety- and stress-related tests in goal-trackers, sign-trackers, and intermediate responders

#### Corticosterone

##### Sample collection

Plasma CORT levels induced by behavioral assays of anxiety-like behavior and physiological restraint were captured using tail nick sampling procedures as described above. Collections took place 24 h before the open field test (time 0, or baseline) and 20, 40, 60, and 80 min post-onset of the test. For restraint-induced CORT profiles, collections took place immediately when rats were placed into the restraining device (time 0, or baseline) and 30 (before being released), 60, 90, and 120 min after the onset of restraint. Rats were transported into the designated room in a staggered fashion, one at a time to begin collections. Repeated nicks were performed on each rat to capture all of the time points. Up to 9 rats remained together in the designated collection room but were on the opposite side of the room from the collection area. Rats were returned to the colony room in a staggered fashion after their last sample was collected.

#### Behavioral testing

##### Elevated plus maze (EPM)

After a ten-day rest period following Experiment 1A, rats were exposed to an elevated plus maze (EPM) test (*refer to experimental timeline*, Figure 1A), considered to be a metric of anxiety-like and risk-taking behavior (Lister, 1987; Walf & Frye, 2007). The apparatus was constructed of four connected arms (each 70 cm from the floor, 45 cm long, and 12 cm wide) made of black Plexiglass and arranged in a cross shape. 45-cm high walls enclosed two opposite arms, and the remaining two were open platforms. A central square (12 x 12 cm) connected all four arms. The test room was dimly lit (40 lux) by a light fixture located above the maze. Prior to the test, rats were transported inside their homecage, along with their cage mate, into the testing room and left undisturbed to acclimate for 30 min. Upon starting the test, each rat was placed in the central platform facing an open arm and allowed to roam freely around the maze for a total of 5 min. The experimenter remained in the room but was distanced from the apparatus in order to be out of the rat’s view. A video-tracking system (Noldus Ethovision 11.5, Leesburg, VA) using a live feed from a digital camera mounted on the ceiling directly above the center of the maze was used to detect and record: 1) latency to enter the open arms for the first time, 2) frequency to enter each arm, and 3) time spent in each arm. Additionally, universally used risk assessment behaviors (RABs, *see* Mikics et al., 2005; Rodgers et al., 1999) were scored manually by the experimenter viewing the live recording. Specifically, the number of times the rat exhibited a bout of grooming, rearing, and protected and unprotected head dips (*i.e*., head dips over the side of the maze while their body was inside an enclosed arm vs. their body being completely exposed on the open platforms) was quantified.

##### Open field test (OFT)

After a ten-day rest period following EPM testing, rats were exposed to an open field test (OFT; *refer to the experimental timeline*, Figure 1A), considered to be another metric of anxiety-like behavior as well as exploratory behavior (Walsh & Cummins, 1976). The OFT test occurred in the same room as the EPM test, and again, paired-housed rats were transported from the colony room to the dimly lit test room and allowed to acclimate for ~30 min before testing began. The open field apparatus was a 4-wall Plexiglass enclosure with an open top and plexiglass floor (100 x 100 x 50 cm). At the start of the test, rats were placed into the same corner (bottom left) of the arena and allowed to roam freely for 5 min. Behavior was video recorded with a digital camera mounted above the apparatus. Noldus Ethovision (11.5, Leesburg, VA) was used to detect: 1) the time spent in the center of the arena (a 50 x 50 cm square drawn in the center), 2) the time spent in the outer edge of the arena (25 cm wide border), 3) the number of entries into the center arena, 4) latency to enter the center of the arena for the first time, and 5) total distance traveled.

##### Restraint

After a ten-day rest period following the OFT, rats underwent a single session of physiological restraint. The restraining device consisted of a white 9 x 12 cm sleeve of flexible Teflon secured with two black Velcro straps attached to a 9 x 3 cm clear Plexiglas platform with a tail slit on one end and breathing holes on the other. Rats were transported in their homecage into the testing room, which was the same as that used for Experiment 1A and OFT time-course measures. Rats were placed into the restrainer and remained there for 30 min.

### Experiment 2: Glucocorticoid receptor (GR) mRNA expression within the hippocampus and prelimbic cortex of goal-trackers, sign-trackers, and intermediate responders

#### Animals

An additional sixty male Sprague-Dawley rats were obtained from Charles River Breeding Labs (C72 and R04) for this experiment. Housing and testing conditions were identical to those described in Experiment 1, except that lights turned on and off at 6:00 and 18:00 h, respectively. Rats were exposed to 2 days of handling prior to behavioral testing, which occurred between 11:00 and 15:00 h.

#### Behavioral testing

##### Pavlovian conditioning approach (PavCA) training

PavCA training and classification of GT, ST, and IR were performed identically to that described above for Experiment 1.

#### Glucocorticoid receptor mRNA expression

##### Tissue collection

Twenty-four hours after completion of the 5^th^ PavCA training session (*refer to the experimental timeline*, Figure 1B), rats underwent live decapitation, and their brains were extracted and immediately flash frozen in 2-methyl butane (−30°C). Brains were stored at −80°C until further processing. Frozen brains were mounted perpendicularly to a metal cryostat chuck using Optimal Cutting Temperature compound (Fisher Healthcare, Thermo Fisher Scientific Kalamazoo, MI, USA) and coated with Shandon M-1 embedding matrix (Thermo Fisher Scientific, Kalamazoo, MI, USA) in preparation for sectioning. Whole brains were coronally sectioned at 10 μm on a cryostat at −20°C. Brain sections were collected, 4.68 to −7.08 mm from Bregma, and directly mounted onto Superfrost Plus microscope slides (Fischer Scientific, Pittsburg, PA, USA), with four sections per slide and ~200 μm between sections on a given slide. Slides were stored at −80°C in preparation for in situ hybridization.

##### Probe synthesis

Probes for in situ hybridization were synthesized in-house using rat mRNA sequences complementary to the RefSeq database number (M14053) for Type II glucocorticoid receptor (GR: insert size 402, insert location nucleotides 765-1167) (*identical to* Garcia-Fuster et al., 2012). All cDNA segments were extracted using a Qiaquick Gel Extraction kit (Qiagen, Valencia, CA), subcloned in Bluescript SK (Stratagene, La Jolla, CA), and confirmed by nucleotide sequencing. The probes were labeled in a reaction mixture of 2μl of linearized DNA specific to the probe, 10X transcription buffer (Epicentre Technologies Madison, WI), 3 μL of S-35-labeled UTP, 10 μL of S-35-labeled ATP, 1 μL of 10 mM CTP and GTP, 1μL of 0.1M dithiothreitol (DTT), 1μl of RNAse inhibitor, and 1μl of T7 RNA polymerase and incubated for 1.5 hours (37°C). Labeled probes were then purified using Micro Bio-Spin 6 Chromatography Column (BioRad, Berkeley, CA), and 1 μl of the probe was counted for subsequent radioactivity dilution calculations. Four to six labelings were used to reach the necessary volume and optimal radioactivity (1-2 million counts per minute/ slide). An additional 1μl of 1M DTT was also added to the labeled mRNA after purification, allowed to incubate at room temperature for 15 min, and stored at −20°C until further use.

##### In situ hybridization

In situ hybridization procedures for hippocampal and prelimbic tissue were performed as described below, independently for each brain region. The radioactive probe was diluted in hybridization buffer (50% formamide, 10% dextran sulfate, 3X saline-sodium citrate buffer, 50 mM sodium phosphate buffer, 1X Denhardt’s solution, 0.1 mg/ml yeast tRNA, and 10 mM DTT) and the volume calculated based on the initial count in order to obtain roughly 1-2 x 10^6^ radioactivity counts per 75 μL of the diluted probe in hybridization buffer. Slide-mounted brain tissue, 4.68 to 2.52 mm (PrL) and −1.08 to −7.08 mm from Bregma (hippocampus), was fixed in 4% paraformaldehyde solution (1 hr), washed in 2X saline-sodium citrate buffer (SSC), and incubated with 0.1 M triethanolamine (TEA) with 0.25% acetic anhydride (10 min). Slides were then dehydrated using ascending ethanol concentrations and air-dried for 1 hour. Hybridization buffer was warmed (37°C) and mixed with the calculated quantity of probe and 1 M DTT (~1% total HB volume). 75 μl of the diluted probe was then applied to coverslips, which were subsequently placed onto the tissue. Slides were then placed in humidity-maintained hybridization chambers soaked with formamide and incubated overnight (~16 hrs) at 55°C. The next day, coverslips were removed, and the slides were rinsed with 2X SSC. Slides were then incubated (1 hr) in RNase A solution (100 μg/mL RNase in Tris buffer with 0.5M NaCl, (37°C), washed in descending concentrations of SSC (2X, 1X, 0.5X), and incubated (1 hr) in 0.1X SSC (65°C). Next, sections were briefly rinsed in H_2_O, dehydrated using ascending ethanol concentrations, and air-dried for 1 hr. Slides were then loaded into film cassettes, separated by regions of interest (*i.e*., PrL and hippocampus) and exposed in a dark room to 35 x 43cm Kodak BioMax MR film (Carestream Health Inc, Rochester, NY, USA) for three weeks for PrL and seven weeks for hippocampal tissue. Extra slides using spare experimental tissue were run concurrently to confirm optimal exposure time. The specificity of the probe was verified using sense strand controls similar to previous studies (Garcia-Fuster et al., 2012; Kabbaj et al., 2000).

##### Quantification

Films were developed using Microtek ScanMaker 1000XL (Fontana, CA, USA) and digitally scanned using SilverFast Lasersoft Imaging software (Sarasota, FL, USA). Signal expression was quantified using ImageJ (National Institutes of Health, Bethesda, MD), a computer-assisted optical densitometry software. For hippocampus GR mRNA quantification, the brush selection tool (size: 15 pixels) was used to trace the curvilinear subregions of interest (CA1, CA2, CA3, and dentate gyrus) throughout the dorsal (−2.64 to −4.56 mm from Bregma) and ventral hippocampus (−4.68 to −6.72 mm from Bregma), using the Rat Brain Atlas (Paxinos and Watson, 2007) for guidance (*see also* Figure 7A). Area (total number of pixels), optical density (darkness of pixels) and integrated optical density (intensity and spread) measurements of the region of interest were taken using a macro that automatically enabled signal above background (3.5 x standard deviation) to be determined. The area (unit, 63 pixels/ 1 mm) and optical density (darkness) were calculated for each of the four hippocampal subregions region across a range of 11-21 sections per rat that spanned the dorsal-ventral gradient of the hippocampus. A single value was calculated for each of the hippocampal subregions per rat, by averaging the values of both hemispheres across multiple sections. Further, given that the dorsal and ventral hippocampus are viewed as neuroanatomically and functionally distinct (*see* Fanselow & Dong, 2010), with the dorsal hippocampus considered to be more involved in cognitive function and the ventral hippocampus in stress and emotion, a single average value was used for dorsal vs. ventral subregions and data were graphed and analyzed separately (*similar to* Romeo et al., 2008).

For PrL GR quantification, the rectangle selection tool (area set to 0.2318 (unit, 63 pixels, 1 mm)) was used across anterior (A)-posterior (P) levels (4.68 to 2.52 mm from Bregma). Optical density was calculated across a range of 3-10 sections per rat that spanned the different A-P levels of the PrL. A single value was also calculated per rat, by averaging the values of both hemispheres across multiple sections. For both hippocampus and PrL quantification, sections with damaged tissue or artifacts that distorted the region of interest were omitted from analyses. During quantification, the experimenter was blind to the phenotype assignments, and the same experimenter quantified both the hippocampus and PrL.

### Experiments 1 and 2: Statistical analysis

Behavioral outcome measures (*i.e*., PavCA, EPM, OFT), plasma corticosterone concentrations, and in situ hybridization measures (*i.e*., area and optical density) were analyzed using the Statistical Package for the Social Sciences (SPSS) program version 24.0 (IBM, Armok, NY, USA). Linear mixed-effects models were performed for PavCA behavior and neuroendocrine measures (corticosterone and GR mRNA levels), using the best fit covariance structure with the lowest Akaike’s information criterion for each set of data. Univariate analysis of variance was performed for behavior exhibited during the EPM and OFT and normality was tested using the Shapiro-Wilk test. When dependent variables failed to meet normality, log 10 or square root transformations were conducted, or a Kruskal-Wallis nonparametric test was performed (using StatView, version 5.0, SAS Institute Inc., Cary, NC, USA). Pearson correlations were performed to determine if there was a significant relationship between baseline CORT levels (pre-vs. post-PavCA) and baseline CORT levels and PavCA behavior. Using univariate analysis of variance, the effect of phenotype was assessed with the predicting variable (*e.g*., change in baseline CORT) set as a covariate. Thus, a significant interaction between the two variables would indicate that the correlational relationship differs between phenotypes. Statistical significance was set at p<0.05, and Bonferroni post-hoc analyses were conducted when significant interactions were detected. All figures were made using GraphPad Prism 7.

## RESULTS

### Experiment 1: Pavlovian conditioned approach behavior and baseline plasma corticosterone profiles

#### PavCA behavior

The following lever-directed (sign-tracking) and food cup-directed (goal-tracking) behaviors were assessed across five consecutive PavCA training sessions and compared between GTs (n=11), IRs (n=17), and STs (n=32): the probability to approach, the number of contacts, and the latency to approach the lever or food-cup during the presentation of the lever-CS (Figure 2). Main Effects of Phenotype, Session, and Phenotype x Session interactions for all behavioral measures are reported in Table 1 (top). There was a significant Effect of Phenotype and Session for all behavioral measures. As expected, STs showed a significantly greater probability to approach the lever (Figure 2A), a greater number of lever contacts (Figure 2C), and shorter latency to approach the lever (Figure 2E), relative to IRs and GTs. These differences in lever-directed behaviors were apparent by the 2^nd^ PavCA training session (Figure 2, *also* Table 2 (top left)). In contrast, relative to STs and IRs, GTs showed a significantly greater probability of approaching the food-cup (Figure 2B), a greater number of food-cup entries (Figure 2D), and a shorter latency to enter the food-cup (Figure 2F). These differences in food cup-directed behavior became apparent by the 3^rd^ PavCA training session (Figure 2, *also* Table 2 (top right)).

**Figure 2.**
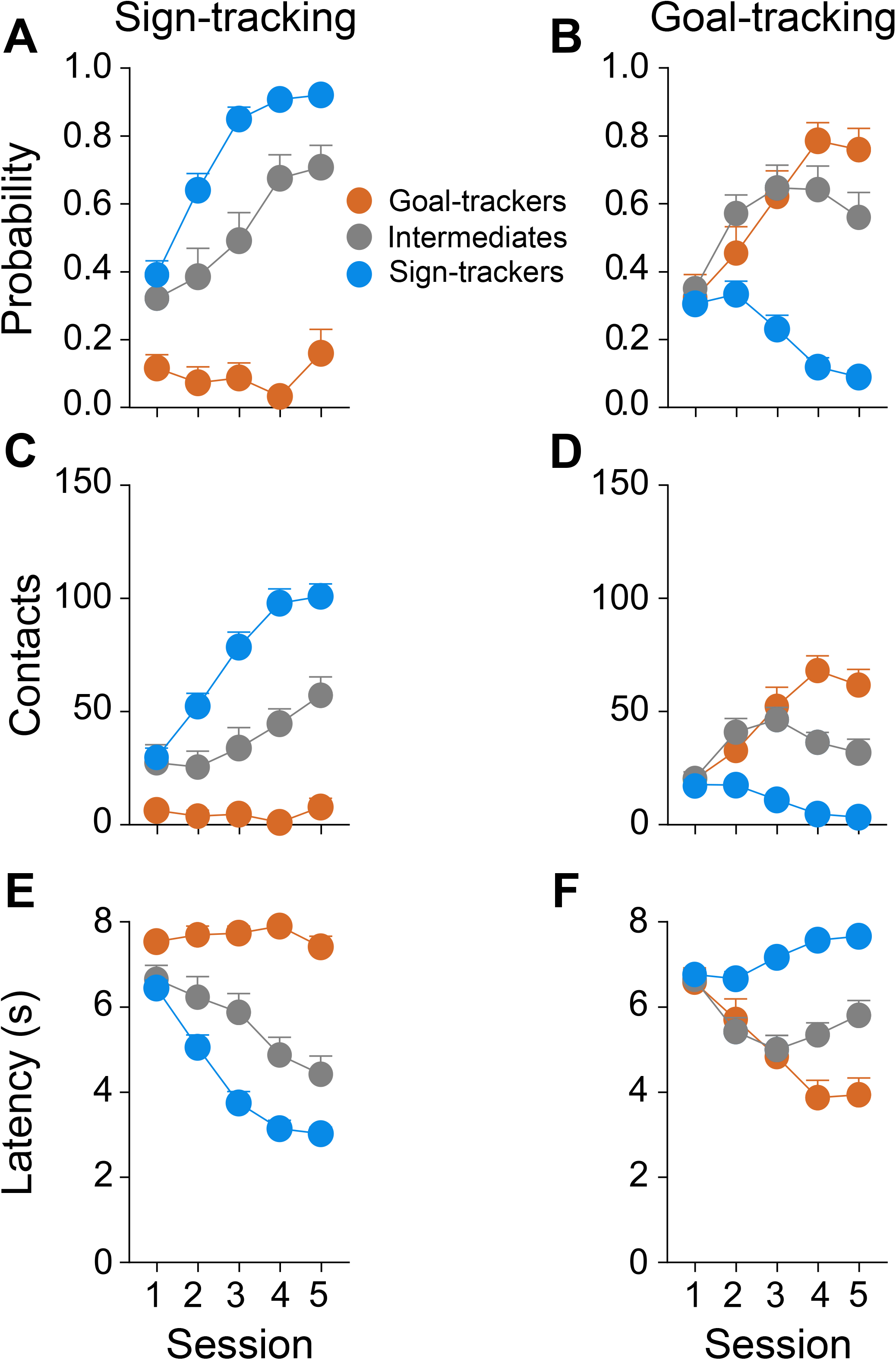
Acquisition of sign-tracking and goal-tracking behavior. Sign-tracking (*i.e.*, lever-directed, left panels) and goal-tracking (*i.e.*, food-cup directed, right panels) behavioral measures were assessed across 5 PavCA sessions. Mean + SEM for probability to: **A**) contact the lever or **B**) enter the food-cup, total number of contacts with **C**) the lever or **D**) the food-cup, and latency to **E**) contact the lever or **F**) enter the food-cup. Rats with a sign-tracking conditioned response were classified as STs (n= 32), those with a goal-tracking conditioned response as GTs (n= 11), and those that vacillated between the two conditioned responses as IRs (n= 17).

**Table 1.** Results from Linear Mixed model analysis for sign-tracking (*i.e.*, lever-directed) and goal-tracking (*i.e.*, food-cup-directed) behaviors. Effect of Phenotype, Session, and Phenotype x Session interactions were analyzed for Experiment 1 (top) and Experiment 2 (bottom). Abbreviations: df1, degrees of freedom numerator, df2, degrees of freedom denominator.

Sign-tracking
|  | Lever Contacts |  |  | Lever Contact Probability |  |  | Lever Contact Latency |  |  |
| --- | --- | --- | --- | --- | --- | --- | --- | --- | --- |
|  | df<sub>1</sub>,df<sub>2</sub> | F-Value | P-Value | df<sub>1</sub>,df<sub>2</sub> | F-Value | P-Value | df<sub>1</sub>,df<sub>2</sub> | F-Value | P-Value |
| Effect of Phenotype | 2, 54.615 | 38.121 | p<0.001 | 2, 57 | 56.803 | p<0.001 | 2, 55.352 | 40.450 | p<0.001 |
| Effect of Session | 4, 87.711 | 15.929 | p<0.001 | 4, 57 | 23.654 | p<0.001 | 4, 79.678 | 35.451 | p<0.001 |
| Phenotype*Session | 8, 87.711 | 8.640 | p<0.001 | 8, 57 | 9.027 | p<0.001 | 8, 79.678 | 11.9.28 | p<0.001 |

Goal-tracking
|  | Food cup Entries |  |  | Food cup Entry Probability |  |  | Food cup Entry Latency |  |  |
| --- | --- | --- | --- | --- | --- | --- | --- | --- | --- |
|  | df<sub>1</sub>,df<sub>2</sub> | F-Value | P-Value | df<sub>1</sub>,df<sub>2</sub> | F-Value | P-Value | df<sub>1</sub>,df<sub>2</sub> | F-Value | P-Value |
| Effect of Phenotype | 2, 63.287 | 52.264 | p<0.001 | 2, 63.003 | 39.844 | p<0.001 | 2, 60.931 | 45.807 | p<0.001 |
| Effect of Session | 4, 96.698 | 14.610 | p<0.001 | 4, 155.487 | 10.960 | p<0.001 | 4, 81.979 | 14.380 | p<0.001 |
| Phenotype*Session | 8, 96.698 | 16.953 | p<0.001 | 8, 155.87 | 12.193 | p<0.001 | 8, 81.979 | 17.313 | p<0.001 |

Experiment 2 Sign-tracking
|  | Lever Contacts |  |  | Lever Contact Probability |  |  | Lever Contact Latency |  |  |
| --- | --- | --- | --- | --- | --- | --- | --- | --- | --- |
|  | df<sub>1</sub>,df<sub>2</sub> | F-Value | P-Value | df<sub>1</sub>,df<sub>2</sub> | F-Value | P-Value | df<sub>1</sub>,df<sub>2</sub> | F-Value | P-Value |
| Effect of Phenotype | 2, 25.738 | 73.590 | p<0.001 | 2,27.026 | 111.836 | p<0.001 | 2, 27.037 | 65.914 | p<0.001 |
| Effect of Session | 4, 30.902 | 17.660 | p<0.001 | 4, 25.527 | 37.054 | p<0.001 | 4, 23.863 | 29.661 | p<0.001 |
| Phenotype*Session | 8, 30.920 | 10.840 | p<0.001 | 8, 25.527 | 17.592 | p<0.001 | 8, 23.885 | 12.406 | p<0.001 |

Goal-tracking
|  | Food cup Entries |  |  | Food cup Entry Probability |  |  | Food cup Entry Latency |  |  |
| --- | --- | --- | --- | --- | --- | --- | --- | --- | --- |
|  | df<sub>1</sub>,df<sub>2</sub> | F-Value | P-Value | df<sub>1</sub>,df<sub>2</sub> | F-Value | P-Value | df<sub>1</sub>,df<sub>2</sub> | F-Value | P-Value |
| Effect of Phenotype | 2, 31.080 | 39.402 | p<0.001 | 2, 27.941 | 37.563 | p<0.001 | 2, 26.445 | 41.540 | p<0.001 |
| Effect of Session | 4, 39.514 | 5.3831 | p= 0.002 | 4, 47.772 | 6.000 | p= 0.001 | 4, 56.426 | 9.495 | p<0.001 |
| Phenotype*Session | 8, 39.631 | 7.976 | p<0.001 | 8, 47.731 | 10.586 | p<0.001 | 8,56.453 | 12.399 | p<0.001 |

**Table 2.** Bonferroni posthoc comparisons between phenotypes for each PavCA session. Signtracking (*i.e.*, lever-directed) and goal-tracking (*i.e*. food-cup-directed) behaviors are included for Experiment 1 (top) and Experiment 2 (bottom). Abbreviations: GTs, goal-tackers, STs, signtrackers, IR, intermediate responders. * p<0.005

Phenotype Comparisons Experiment 1
|  | Sign-tracking<br>Lever Contacts |  |  |  |  | Goal-tracking<br>Food cup Entries |  |  |  |  |
| --- | --- | --- | --- | --- | --- | --- | --- | --- | --- | --- |
|  | 1 | 2 | 3 | 4 | 5 | 1 | 2 | 3 | 4 | 5 |
| GT vs. IR | p= 0.090 | p= 0.152 | p= 0.083 | p= 0.001* | p<0.001* | p= 1.000 | p= 0.820 | p= 1.000 | p<0.001* | p<0.001* |
| GT vs. ST | p= 0.024* | p<0.001* | p<0.001* | p<0.001* | p<0.001* | p= 1.000 | p= 0.072 | p<0.001* | p<0.001* | p<0.001* |
| ST vs. IR | p= 1.000 | p= 0.007* | p<0.001* | p<0.001* | p<0.001* | p= 1.000 | p<0.001* | p<0.001* | p<0.001* | p<0.001* |
| Lever Contact Probability |  |  |  |  | Food cup Entry Probability |  |  |  |  |  |
|  | 1 | 2 | 3 | 4 | 5 | 1 | 2 | 3 | 4 | 5 |
| GT vs. IR | p= 0.074 | p= 0.018* | p<0.001* | p<0.001* | p<0.001* | p= 1.000 | p= 0.542 | p= 1.000 | p= 0.226 | p= 0.051 |
| GT vs. ST | p= 0.004* | p<0.001* | p<0.001* | p<0.001* | p<0.001* | p= 1.000 | p= 0.382 | p<0.001* | p<0.001* | p<0.001* |
| ST vs. IR | p=0.930 | p= 0.012* | p<0.001* | p<0.001* | p= 0.001* | p= 1.000 | p= 0.002* | p<0.001* | p<0.001* | p<0.001* |
| Lever Contact Latency |  |  |  |  | Food cup Entry Latency |  |  |  |  |  |
|  | 1 | 2 | 3 | 4 | 5 | 1 | 2 | 3 | 4 | 5 |
| GT vs. IR | p= 0.092 | p= 0.057 | p= 0.006* | p<0.001* | p<0.001* | p= 1.000 | p= 1.00 | p<0.001* | p<0.001* | p<0.001* |
| GT vs. ST | p= 0.011* | p<0.001* | p<0.001* | p<0.001* | p<0.001* | p= 1.000 | p= 0.081 | p<0.001* | p<0.001* | p<0.001* |
| ST vs. IR | p= 1.000 | p= 0.047* | p<0.001* | p<0.001* | p= 0.001* | p= 1.000 | p= 0.003* | p<0.001* | p<0.001* | p<0.001* |

Experiment 2
|  | Sign-tracking<br>Lever Contacts |  |  |  |  | Goal-tracking<br>Food cup Entries |  |  |  |  |
| --- | --- | --- | --- | --- | --- | --- | --- | --- | --- | --- |
|  | 1 | 2 | 3 | 4 | 5 | 1 | 2 | 3 | 4 | 5 |
| GT vs. IR | p= 0.852 | p= 0.005* | p= 0.016* | p= 0.001* | p<0.001* | p=0.959 | p=1.000 | p= 0.063 | p<0.001* | p<0.001* |
| GT vs. ST | p= 0.001* | p<0.001* | p<0.001* | p<0.001* | p<0.001* | p= 0.073 | p= 0.052 | p<0.001* | p<0.001* | p<0.001* |
| ST vs. IR | p= 0.010* | p<0.001* | p<0.001* | p<0.001* | p<0.001* | p= 0.547 | p=0.217 | p= 0.001* | p<0.001* | p= 0.003* |
| Lever Contact Probability |  |  |  |  | Food cup Entry Probability |  |  |  |  |  |
|  | 1 | 2 | 3 | 4 | 5 | 1 | 2 | 3 | 4 | 5 |
| GT vs. IR | p= 0.211 | p<0.001* | p<0.001* | p<0.001* | p<0.001* | p= 1.000 | p=1.000 | p= 0.820 | p= 0.093 | p= 0.005* |
| GT vs. ST | p<0.001* | p<0.001* | p<0.001* | p<0.001* | p<0.001* | p= 0.163 | p= 0.138 | p<0.001* | p<0.001* | p<0.001* |
| ST vs. IR | p=0.018* | p<0.001* | p= 0.004* | p= 0.003* | p= 0.003* | p= 0.331 | p= 0.359 | p<0.001* | p<0.001* | p<0.001* |
| Lever Contact Latency |  |  |  |  | Food cup Entry Latency |  |  |  |  |  |
|  | 1 | 2 | 3 | 4 | 5 | 1 | 2 | 3 | 4 | 5 |
| GT vs. IR | p= 0.649 | p= 0.005* | p<0.001* | p<0.001* | p<0.001* | p= 1.000 | p= 1.00 | p= 0.072 | p<0.001* | p<0.001* |
| GT vs. ST | p<0.001* | p<0.001* | p<0.001* | p<0.001* | p<0.001* | p= 0.285 | p= 0.096 | p<0.001* | p<0.001* | p<0.001* |
| ST vs. IR | p= 0.013* | p= 0.001* | p<0.001* | p<0.001* | p= 0.002* | p= 0.420 | p= 0.408 | p= 0.006* | p<0.001* | p<0.001* |

#### Baseline CORT levels pre- and post-PavCA

##### CORT levels

Overall, pre- and post-PavCA baseline plasma CORT levels did not significantly differ between Phenotypes (GTs n=10, IRs n=17, STs n=32) [Effect of Phenotype: F_(2,57.691)_=2.325, *p*=0.107] (Figure 3). Relative to pre-PavCA, post-PavCA baseline CORT levels were significantly higher [Effect of Timepoint: F_(1,53.246)_=20.180, *p*<0.001], rising from an overall average of 56 ng/mL (pre-PavCA) to 108 ng/mL (post-PavCA). While, baseline CORT levels appear to rise with the experience of PavCA training, the extent to which CORT increased was not dependent on Phenotype [Time-point x Phenotype interaction: F_(2, 52.633)_=0.535, *p*=0.589]. These data are in agreement with prior studies (Flagel et al., 2009; Tomie et al., 2000), demonstrating that pre-PavCA baseline plasma CORT levels do not significantly differ between Phenotypes, and extend these findings to show that baseline plasma CORT levels also do not differ between Phenotypes after the development of a conditioned response.

**Figure 3.**
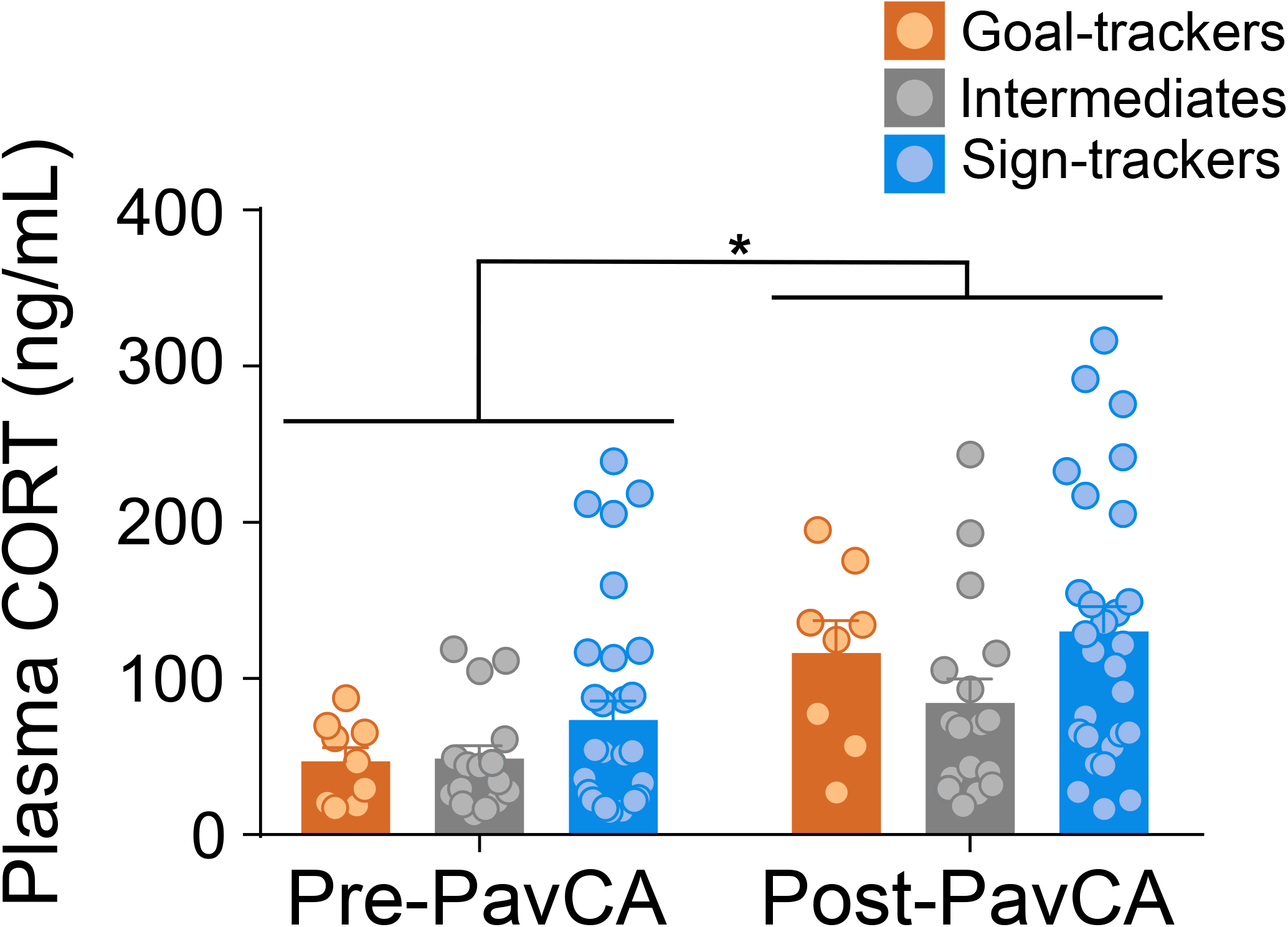
“Baseline” corticosterone levels before and after Pavlovian conditioned approach training. Mean + SEM for baseline plasma CORT levels prior to (Pre-PavCA) and following PavCA training experience (Post-PavCA). Basal plasma CORT levels increased with Pavlovian training experience (*, p= 0.001; n=60; GT n=10, IR=17, ST=32).

##### Correlations

To further investigate the relationship between baseline CORT levels and cue-motivated behavior, we performed correlational analyses. Pre-PavCA baseline levels significantly correlated with post-PavCA baseline levels [r= 0.449, *p*=0.001]; but pre-PavCA baseline levels did not correlate with the behavioral phenotype that emerges with PavCA training (*i.e*., the average PavCA index from session 4 and 5) [r= 0.198, *p*=0.143]. In relation, there was not a significant correlation between the change in baseline CORT levels from pre-to post-PavCA (*i.e*., Δ CORT) and the magnitude of change in the conditioned response from the onset of training (Session 1) to the end of training (Session 5) (*i.e*., Δ PavCA index) [r= 0.121, *p*=0.402, Table 3]. While these data provide little evidence of a relationship between baseline CORT levels and the propensity to attribute incentive salience to a reward cue, prior findings (*e.g*., Flagel et al., 2009) prompted us to further assess this relationship within each phenotype group. When analyzed independently, there is a significant positive correlation between Δ CORT and Δ PavCA index for STs [r=0.470, *p*=0.013], and a non-significant negative correlation in GTs [r=-0.301, *p*= 0.512] and IRs [r=-0.058, *p*=0.830]. Notably, these results should be interpreted with caution as there was not an interaction with the predicting variable [Phenotype x Δ CORT: F2,4=1.858, *p*=0.168] to suggest that this relationship significantly differs between phenotypes, and the sample size for GTs is quite low. Nonetheless, it should also be noted that similar patterns held true for other measures, with significant correlations between baseline CORT values and behavioral measures for STs, but not for GTs or IRs (*see* Table 3).

**Table 3.**
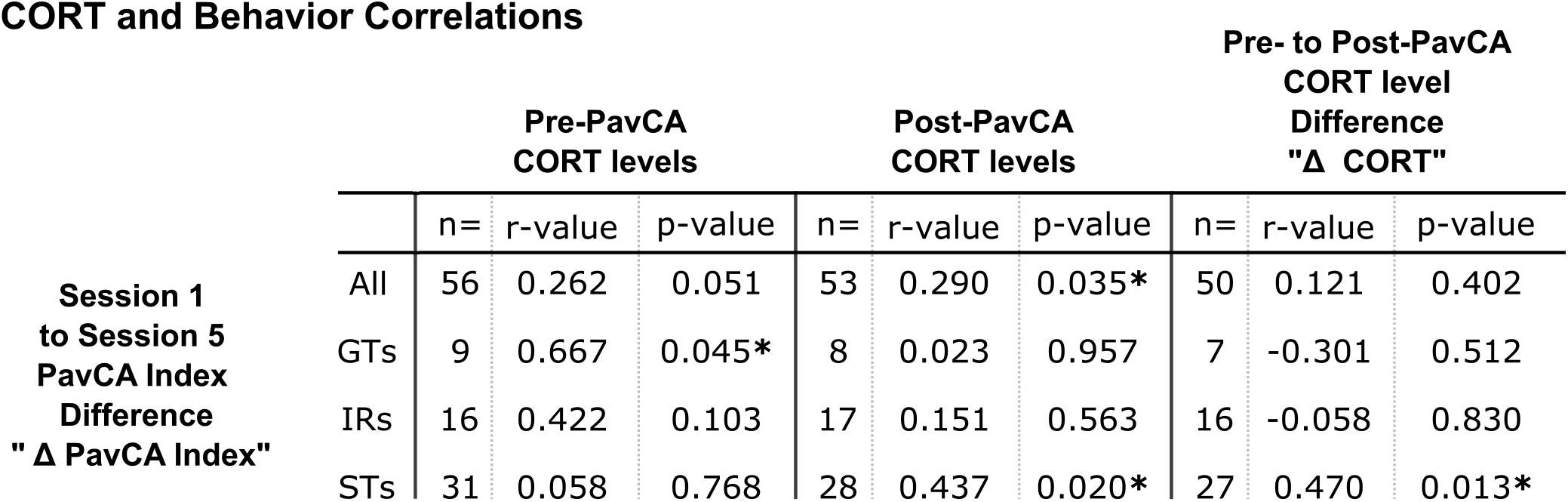
Results of Pearson correlation analysis between the change in the PavCA Index from Session 1 to Session 5 (Δ PavCA Index) and baseline CORT profiles (Pre-PavCA, Post-PavCA, and the change in baseline values, Δ CORT) for the entire population (*i.e.*, All) and for each phenotype separately. * p<0.005

### Experiment 1B: Behavioral and corticosterone response to anxiety- and stress-related tests in goal-trackers, sign-trackers, and intermediate responders

#### Elevated Plus Maze

GTs (n=11) and STs (n=14) did not significantly differ on any behavioral outcome measure of the EPM test, but statistical analysis revealed significant differences relative to their intermediate responder counterparts (n=13). While all rats spent the most time [Effect of Zone: F_(2,105)_=140.397, *p*<0.001] inside the closed arms 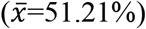, relative to the open arms 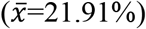 or center square 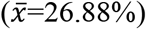, IRs spent significantly less time [Phenotype x Zone interaction: F_(4,105)_=2.762, *p*=0.031] inside the open arms, relative to GTs (*p*=0.011) (Figure 4B). However, the latency to enter the open arms for the first time was similar across all Phenotypes [Effect of Phenotype: F_(2,35)_=0.187, *p*=0.83] (data not shown) and, in general, relative to IRs, the extreme Phenotypes (GTs, *p*=0.02, and STs, *p*=0.049) entered different zones of the EPM more frequently [Effect of Phenotype: F_(2,105)_=6.744, *p*=0.002] (data not shown). Additionally, there were no significant differences between Phenotypes for any of the risk assessment behaviors during the EPM test: frequency of grooming [Kruskal-Wallis test, Effect of Phenotype: χ^2^_(2)_=1.984, *p*=0.371], rearing [Effect of Phenotype: F_(2,35)_=2.232, *p*= 0.122], protected head dips [Effect of Phenotype: F_(2,35)_=0.496, *p*=0.613], or unprotected head dips [Effect of Phenotype: F_(2,24)_=0.207, *p*=0.814] (data not shown).

**Figure 4.**
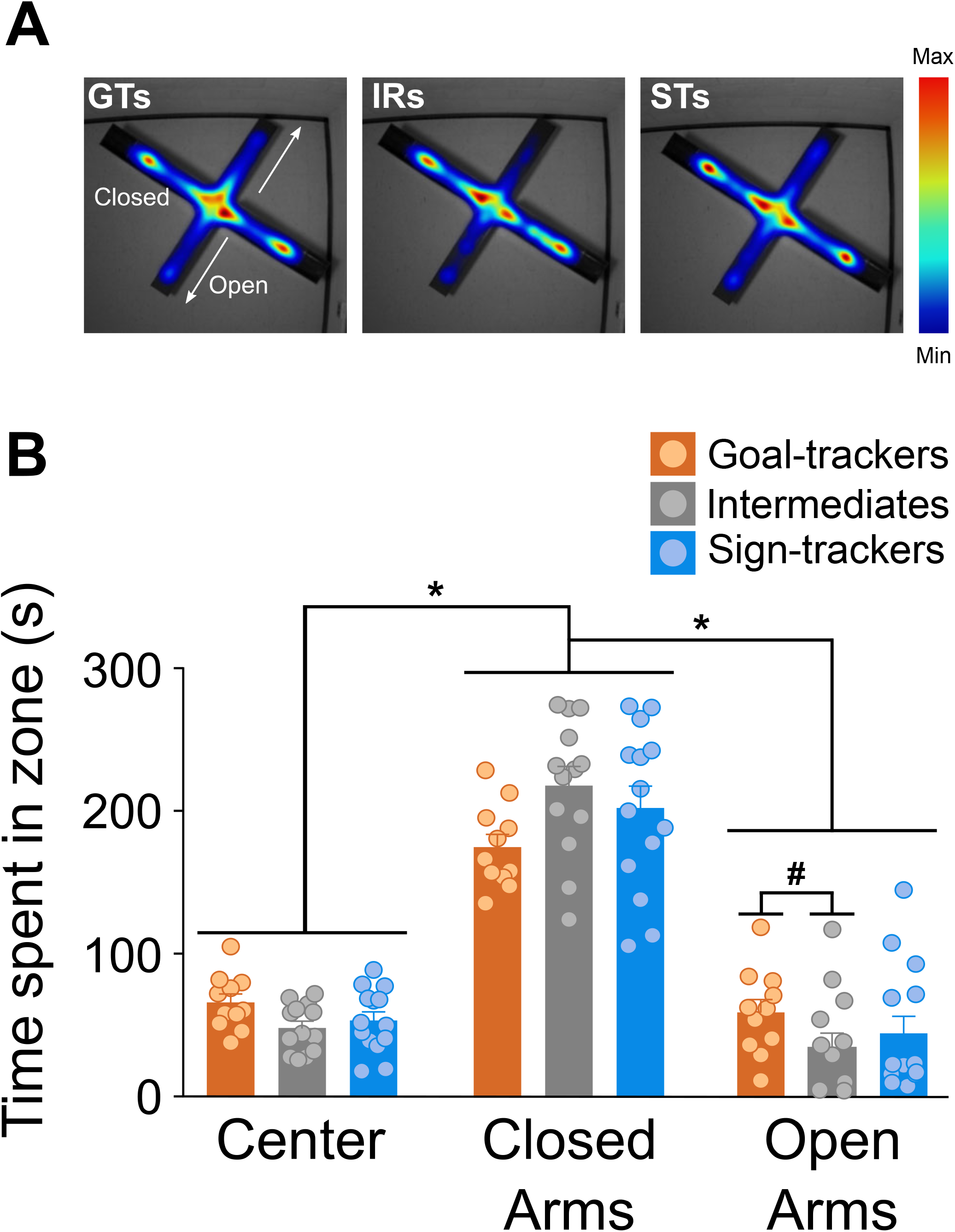
Elevated plus maze. **A)** Heat map representations for the average time spent in each zone during the 5-min EPM test for each phenotype. **B**) Mean + SEM for the time spent in each zone of the elevated plus maze for goal-trackers (n=11), intermediate responders (n=13), and sign-trackers (n=14). All rats spent significantly more time in the closed arms compared to the open arms and center of the maze (*, p<.001). There was not a significant difference between GTs and STs in the amount of time spent in either the center of the arena or the open or closed arms. IRs spent significantly less time in the open arms, relative to GTs (#, p<0.05).

#### Open field test

There were no significant differences between Phenotypes in their behavior on the OFT. All rats spent a comparable amount of time in the outer edge of the arena [Kruskal-Wallis, Effect of Phenotype: χ^2^_(2)_=2 .012, *p*=0.366], with little time spent in the center of the arena (Figure 5). There were no significant differences in the number of entries to the center of the arena [Kruskal-Wallis Effect of Phenotype: χ^2^_(2)_=3 .029, *p*=0.220] (data not shown), latency to enter the center of the arena [Kruskal-Wallis Effect of Phenotype: χ^2^_(2)_=2 .345, *p*=0.310] (data not shown), or time spent in the center of the arena [Kruskal-Wallis, Effect of Phenotype: χ^2^_(2)_=2 .053, *p*=0.358] (Figure 5). The distance traveled during the OFT was also similar between phenotypes [Kruskal-Wallis Effect of Phenotype: χ^2^_(2)_=3.287, *p*=0.193] (data not shown). These data suggest that STs, GTs and IRs do not differ in anxiety-like behavior, which is consistent with the data described above from the EPM.

**Figure 5.**
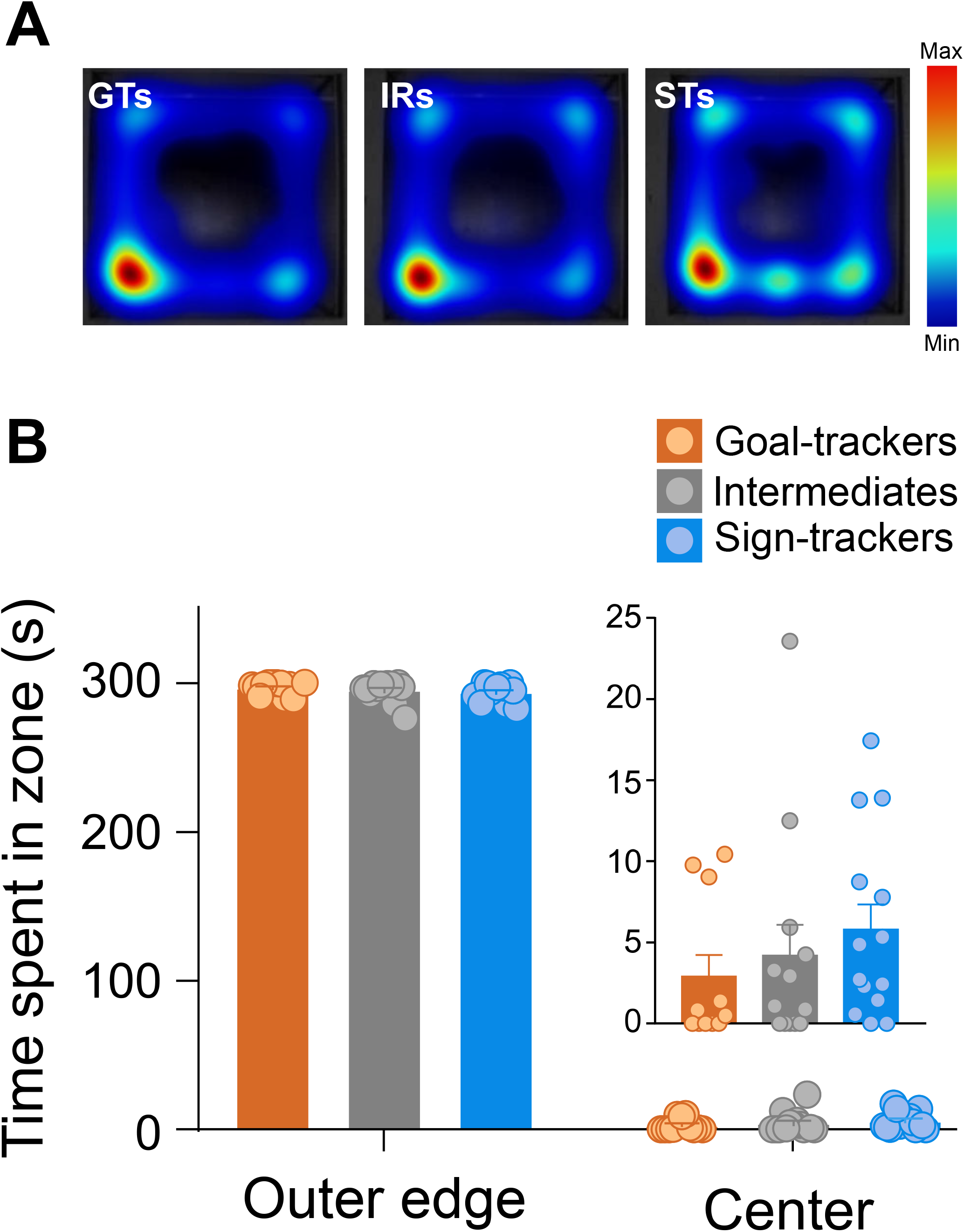
Open field test. **A**) Heat map representations for the average time spent in each zone (outer edge vs. center) during the 5-min OFT for each phenotype. **B**) Mean + SEM for time spent in the outer edge or center of the arena for goal-trackers (n=11), intermediate responders (n=13), and sign-trackers (n=14). All rats spent significantly more time on the outer edge of the arena compared to the center. Time spent in the center of the arena is shown as an inset on a different scale for illustration purposes. There was not a significant difference between phenotypes for the amount of time spent in the center of the arena.

#### Corticosterone response

##### Corticosterone response to OFT

Exposure to the OFT elicited a CORT response [Effect of Time: F_(4,44.652)_=12.849, *p*<0.001], with a significant rise relative to baseline at 20, 40, 60, and 80 min post-OFT onset. Although the CORT response was decreased at 80-min relative to the peak response (40 vs. 80 min, *p*<0.001), a return to baseline levels was not captured with this time course (baseline vs. 80 min, *p*= 0.041). Nonetheless, the CORT response to the OFT did not significantly differ between phenotypes [Effect of Phenotype: F_(2, 35.564)_=0.215, *p*=0.808; Time x Phenotype interaction: F_(8,45.180)_=.718, *p*=.675)] (Figure 6A).

**Figure 6.**
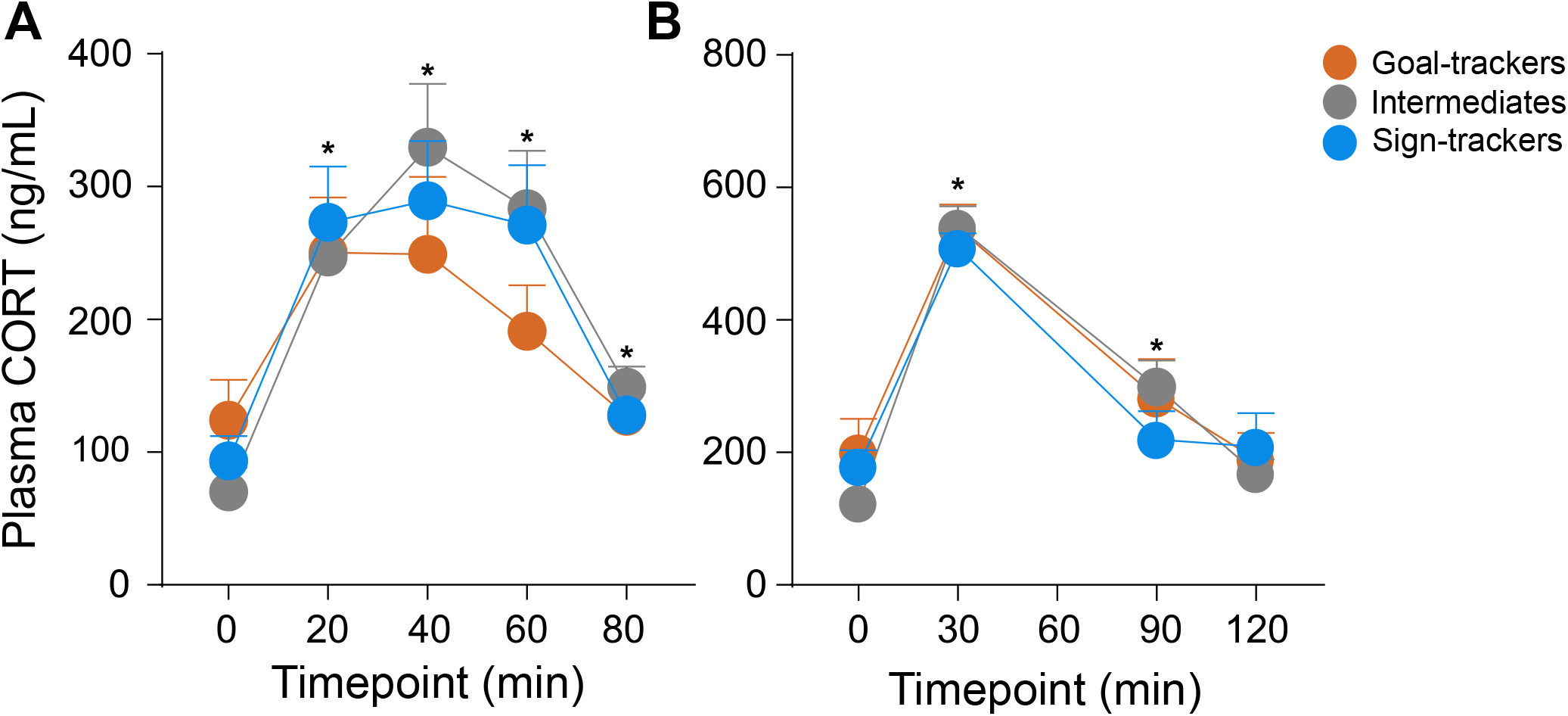
Corticosterone response to the open field test and acute physiological restraint. **A)** Mean + SEM for plasma CORT levels 0,20,40,60, and 80-mins post-onset of the OFT for goal-trackers (n=11), intermediate responders (n=13), and sign-trackers (n=14). There was a significant increase in CORT induced by the OFT at 20, 40, 60, and 80-min time-points (*, p<0.001), but no significant difference between phenotypes. **B**) Mean + SEM for plasma CORT levels 0,30,90, and120-mins post-onset of acute restraint for goal-trackers (n=11), intermediate responders (n=13) and sign-trackers (n=12). There was a significant increase in CORT induced by restraint at 30 and 90-min time-points (*, p<.001), but no significant difference between phenotypes.

**Figure 7.**
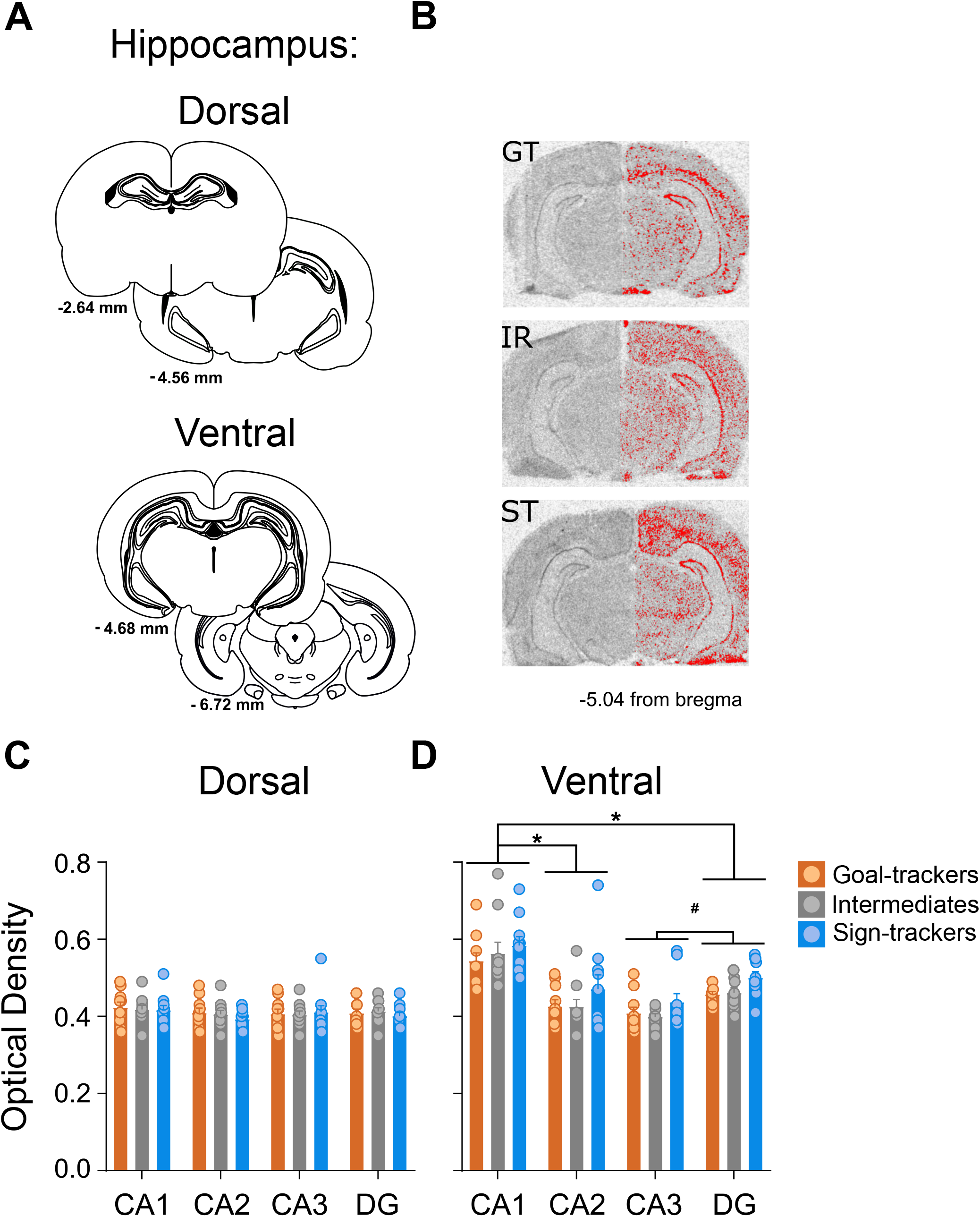
Glucocorticoid receptor mRNA expression in the dorsal and ventral hippocampus. **A)** Coronal brain sections representing bregma coordinates used to quantify glucocorticoid receptor (GR) mRNA expression (*Adapted from* Paxinos and Watson, 2007). **B**) Representative in situ images for a GT, IR, and ST rat with signal above threshold represented (in red) on the right hemisphere. **C-D**) Mean + SEM optical density for GR mRNA in subregions of the **C)** dorsal and **D)** ventral hippocampus for goal-trackers (n=10), intermediate responders (n=10) and sign-trackers (n=10). In the ventral hippocampus, GR mRNA varied between subregions (*, p<0.001 vs. CA1, #, p<0.001 vs. DG). Relative to goal-trackers and intermediate responders, sign-trackers show greater GR mRNA density across subregions.

##### Corticosterone response to physiological restraint

Acute physiological restraint (30 min) elicited a CORT response [Effect of Time: F_(3,28.058)_=157.308, *p*<0.001], with a significant rise relative to baseline at 30 and 90 min, and return to baseline levels at 120 min post-onset of restraint. There was not a significant difference in the CORT response to acute restraint between Phenotypes [Effect of Phenotype: F_(2,32.084)_=.114, *p*=0.893; Time x Phenotype interaction: F_(6,29.646)_=1.568, *p*=0.191] (Figure 6B).

### Experiment 2: Glucocorticoid receptor (GR) mRNA expression within the hippocampus and prelimbic cortex of goal-trackers, sign-trackers, and intermediate responders

#### PavCA behavior

Similar to Experiment 1, there were significant Effects of Phenotype, Session, and Phenotype x Session interactions for all behavioral measures reported in Table 1 (bottom) (data are not shown in graphical format). Differences between Phenotypes were apparent for lever- and food cup-directed behavior as early as the first PavCA training session (*see* Table 2 (bottom)).

#### GR mRNA expression

##### Dorsal hippocampus

There were no significant differences between Phenotypes in GR mRNA expression (*i.e*., optical density) in the dorsal hippocampus [Effect of Phenotype: F_(2,108)_=0.233, *p*=0.793], and no significant difference in expression patterns between subregions of the dorsal hippocampus [Effect of Subregion: F_(3,108)_=1.089, *p*=0.357] (Figure 7C). Given the anatomical variability in size between subregions (*see schematic* Figure 7A), significant differences in area were detected [Effect of Subregion: F_(3, 108)_=1020.291, *p*<0.001] (data not shown); CA1 subregion contained the largest area 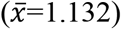, while CA2 the smallest 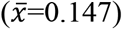. However, the regions of interest were manually outlined (CA1, CA2, CA3, DG), and the area was not dependent on Phenotype [Effect of Phenotype: F_(2,108)_=0.118, *p*=0.889; Phenotype x Subregion interaction: F_(6,108)_=0.417, *p*=0.866], indicating that the selection of regions of interest was consistent across phenotypes.

##### Ventral hippocampus

Unlike the dorsal hippocampus, GR mRNA expression significantly differed between phenotypes [Effect of Phenotype: F_(2, 108)_=4.601, *p*=0.012] and Subregions [Effect of Subregion: F_(3,108)_=30.464, p<0.001] in the ventral hippocampus. STs 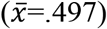 had greater optical density relative to GTs (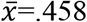, *p*=0.022) and IRs (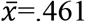, *p*=0.040), and there was greater optical density in CA1 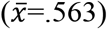 relative to CA2 (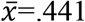, *p*<0.001), CA3 (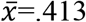, *p*<0.001), and DG (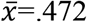, *p*<0.001) (Figure 7D). In addition, optical density in DG was greater than that in CA3 (*p*=0.004). Like the dorsal hippocampus, area was significantly different between Subregions [Subregion: F_(3,108)_=172.935, *p*<0.001], but not between Phenotypes [Effect of Phenotype: F_(2,108)_=0.579, *p*=0.562; Phenotype x Subregion interaction: F_(6,108)_=0.876, *p*=0.515] (data not shown).

##### Prelimbic cortex

Given GR mRNA expression within the PrL did not differ between anterior-posterior levels [Effect of AP Level: F _6, 146_ =0.191, *p*= 0.979; AP Level x Phenotype interaction: F_12, 146_=0.298, *p*=0.989], GR mRNA values were averaged per rat across all AP levels. GR mRNA expression did not differ between phenotypes in the prelimbic cortex [Effect of Phenotype: F_(2,26)_=0.811, *p*=0.455; Figure 8C].

**Figure 8.**
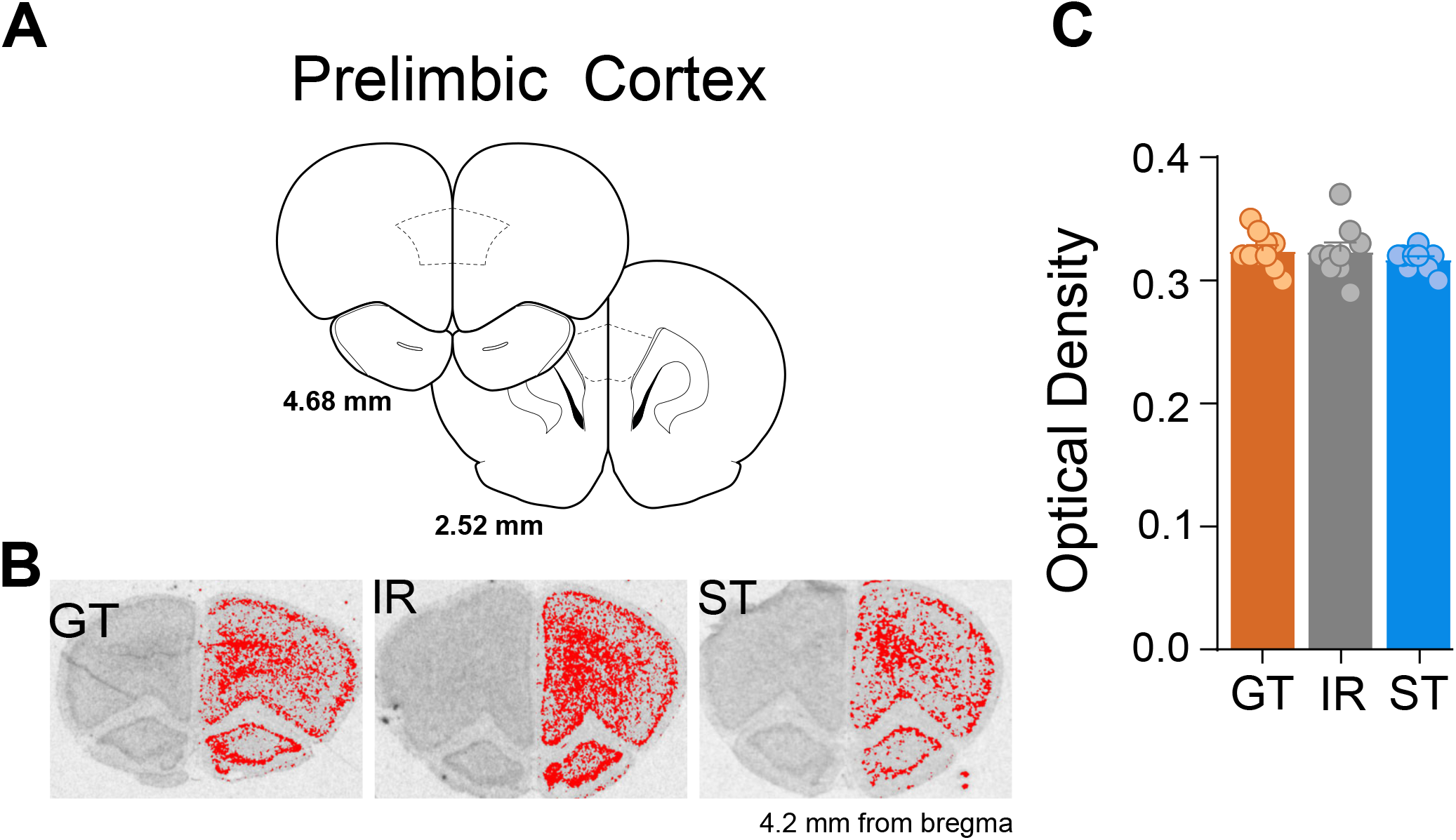
Glucocorticoid receptor mRNA expression in the prelimbic cortex. **A)** Coronal brain sections representing bregma coordinates used to quantify glucocorticoid receptor (GR) mRNA expression (*Adapted from* Paxinos and Watson, 2007). **B**) Representative in situ images for a GT, IR, and ST rat with signal above threshold represented (in red) on the right hemisphere. **C**) Mean + SEM optical density for GR mRNA in the prelimbic cortex (PrL) for goal-trackers (n=10), intermediate responders (n=10) and sign-trackers (n=10).

## Discussion

The present studies examined whether differences in behavioral and neuroendocrine measures of stress and anxiety are included amongst the co-existing traits associated with the propensity to attribute incentive salience to reward cues. We report three main findings. First, there is a general increase in baseline plasma CORT levels over the course of associative cue-learning, and this increase is independent of the Pavlovian conditioned response that emerges. Second, behavioral and CORT responses to environmental challenges that elicit aversive arousal do not differ between goal-trackers and sign-trackers. Third, sign-trackers have greater expression of GR mRNA in the ventral hippocampus relative to goal-trackers and intermediate responders, whereas GR mRNA in the prelimbic cortex is comparable between phenotypes.

The basis of our understanding of CORT function in appetitive Pavlovian conditioning stems from the work of Tomie et al. (2000) who demonstrated that cue-food associations elicit an increase in plasma CORT. Subsequently, it was shown that, relative to GTs, STs exhibit a greater rise in plasma CORT following a single Pavlovian conditioning session; that is, before the development of a conditioned response (Flagel et al., 2009). Further, prior to PavCA training, baseline CORT is similar across GTs, IRs, and STs (Flagel et al., 2009). To date, however, baseline plasma CORT had not been systematically assessed in the same rat to determine if CORT profiles change as a consequence of cue-learning. Thus, in Experiment 1A we compared, within the same rat, baseline CORT concentrations at a “naïve” state of learning (pre-PavCA) and once a conditioned response had been acquired (post-PavCA). There was a significant rise in baseline plasma CORT levels with the development of a conditioned response, and pre-PavCA CORT levels correlated with post-PavCA CORT levels. Contrary to our hypothesis, however, this rise in baseline CORT was not dependent on the innate cue-learning strategy that was employed, as levels did not significantly differ between goal-trackers and sign-trackers. Further, baseline CORT levels at a “naïve” state of Pavlovian learning did not predict the behavior that emerged with Pavlovian training, and the change in baseline CORT did not significantly correlate with the change in Pavlovian conditioned approach behavior over the course of learning. When phenotypes were assessed independently, however, the change in baseline CORT levels was significantly correlated with the change in Pavlovian conditioned approach behavior in sign-trackers, and this relationship was not significant in goal-trackers or intermediate responders. While these correlational data do not prompt strong conclusions in and of themselves, they align with prior reports demonstrating a relationship between CORT and the development of sign-tracking behavior (Flagel et al., 2009, Tomie et al., 2000).

One of the primary roles of CORT is to act across the body and brain to broadly mediate the stress response (Herman et al., 2016). Thus, we wanted to determine whether differences in plasma CORT are present in goal-trackers vs. sign-trackers in contexts outside of Pavlovian conditioning and, explicitly, in response to paradigms related to stress and anxiety. As we hypothesized, Experiment 1B showed no differences between phenotypes in CORT response to an open field test or physiological restraint. Further, goal-trackers and sign-trackers did not differ in their behavioral response to the elevated plus maze or open field test. Notably, rats spent little time within the center of the open field arena, which may be indicative of especially aversive conditions that could have precluded group differences. Nonetheless, these findings are consistent with those previously reported by Vanhille et al. (2015), who showed no pre-existing differences in behavior on the elevated plus maze test in rats that were later characterized as sign-trackers or goal-trackers; and those reported by Harb and Almeida (2014) who showed no differences in behavior on the open field test in mice characterized as sign-trackers or goal-trackers. In contrast to the present findings, however, Harb and Almeida (2014) did report a significant difference in peak CORT response following an acute stressor, with sign-tracker mice exhibiting a greater peak relative to goal-trackers or intermediate responders. These discrepant findings are likely due to differences in experimental procedures, including the species used and the nature and intensity of the stressor. In this regard, we note that the repeated testing implemented in the current study may have affected the CORT response in a manner that precluded observable differences (*e.g.* see Dallman et al., 2004). Indeed, it is possible that differences in the CORT profile in response to physiological restraint were not apparent because of a ceiling effect, as both baseline and peak CORT levels were high across all animals. Taken together, given that goal-trackers and sign-trackers behaved similarly on both tests of anxiety-like behavior, and showed no significant differences in CORT response to either the open field test or physiological restraint, we conclude, based on the paradigms implemented here, that individual differences in aversive arousal are not captured by the goal-tracker/sign-tracker animal model.

It is important to note that other indices of aversive arousal, including fear conditioning and the associated freezing response have been reported to differ between goal-trackers and sign-trackers (Morrow et al., 2011; Morrow et al., 2015). Specifically, relative to goal-trackers, sign-trackers are more fearful of discrete cues that predict footshock (Morrow et al., 2011), and show exaggerated incubation of their fear response (Morrow et al., 2015). Yet, goal-trackers exhibit greater contextual fear when placed back into a fear-conditioning context in the absence of discrete cues (Morrow et al., 2011). Thus, these differences seem to be specific to learning the value of the discrete cue, rather than differences in aversive arousal or stress-reactivity. Others have shown that CORT plays a critical role in fear conditioning, beyond the stress component (*e.g*., Marchand et al., 2007; Zorawski & Killcross, 2002), but this has yet to be assessed within the context of the goal-tracker/sign-tracker animal model and will be the focus of future investigations.

While the current findings and those of others (Harb & Almeida, 2014; Vanhille et al., 2015) demonstrate that goal-trackers and sign-trackers respond similarly to behavioral assays indicative of stress and anxiety, exposure to stress has been shown to alter the propensity to attribute incentive salience to reward cues (Fitzpatrick et al., 2019; Hynes et al., 2018; Lomanowska et al., 2011). Rats exposed to stress early in life exhibit greater sign-tracking behavior in adulthood (Hynes et al., 2018; Lomanowska et al., 2011). In contrast, adult rats exposed to a single prolonged stressor show an attenuation of sign-tracking behavior (Fitzpatrick et al., 2019). Thus, the impact of stress on the propensity to sign-track appears to be dependent on the type of stressor and timing of exposure. In light of the current findings, we postulate that the neural processes underlying these reported stress-induced effects (Fitzpatrick et al., 2019; Hynes et al., 2018; Lomanowska et al., 2011) go beyond CORT and the HPA axis, and include components of the cortico-thalamic-striatal “motive” circuit, which is differentially engaged in sign-trackers vs. goal-trackers (Flagel, Cameron, et al., 2011; *see also* Kuhn et al., 2018).

The hippocampus (*e.g.*, Maccari et al., 1991, *for review see* Barr et al., 2017) and prefrontal cortex (*e.g.*, Butts & Phillips, 2013; Deroche-Gamonet et al., 2003) are two brain regions that may serve as a potential neural interface between the stress and the “motive” circuitry. Glucocorticoid receptors are densely expressed within the hippocampus (Reul & de Kloet, 1985) and CORT-GR interactions within this brain region play a critical role in the negative feedback system that acts to maintain homeostatic levels of CORT in the face of physiological or environmental challenges (Herman et al., 2012). Specifically, greater GR mRNA expression in the hippocampus has been associated with more rapid negative feedback, or return to baseline CORT levels (Liberzon et al., 1999; Meaney et al., 1996). In the current study, we did not observe phenotypic differences in circulating CORT levels either under baseline conditions or in response to environmental challenges (*i.e.*, open field test or physiological restraint). Yet, we did find that, relative to goal-trackers and intermediate responders, sign-trackers have significantly greater expression of GR mRNA in the ventral hippocampus. Although we did not hypothesize phenotypic differences in GR expression to be specific to the ventral hippocampus, the fact that these differences are not apparent in the dorsal hippocampus may explain why we did not observe differences in circulating levels of CORT. Indeed, while the dorsal hippocampus has been shown to play a role in stress-induced negative feedback (Feldman & Weidenfeld, 1993, 1999), the ventral hippocampus has been reported to regulate tonic levels of CORT (Herman et al., 1992). Other findings suggest that the engagement of the ventral vs. dorsal hippocampus is dependent on the type of stressor (*e.g.*, Dorey et al., 2012; Herman et al., 2005; Maggio & Segal, 2007). Thus, additional studies are warranted to further investigate the role of GR expression in the ventral hippocampus within the context of the stress response and negative feedback regulation. Nonetheless, we propose that the phenotypic differences reported here in GR expression in the ventral hippocampus are directly related to motivated behavior and reward learning, rather than stress regulation. Further, lesions of the ventral, but not the dorsal, hippocampus decrease the propensity to sign-track (Fitzpatrick et al., 2016). While it is remains to be determined whether these lesion effects are dependent on GR function, it should be noted that systemic administration of a GR antagonist similarly attenuates the acquisition of sign-tracking behavior (Rice et al., 2018; Rice et al., 2019).

Like the hippocampus, the prefrontal cortex has inhibitory effects on HPA axis activity (*for review, see* Herman et al., 2003). Specifically, GRs within the prelimbic (PrL) subregion of the prefrontal cortex are implicated in response to acute stressors, with their absence increasing the CORT response (McKlveen et al., 2013). GRs within the PrL also influence reward-related mediators, like dopamine (Butts & Phillips, 2013). Within the context of the GT/ST model, the prelimbic cortex is recognized as an integral component of “top-down” control over incentive salience attribution (*e.g.*, Campus et al., 2019); with sign-trackers deficient in cortical control, relative to goal-trackers (*for review, see* Sarter & Phillips, 2018). Given that GTs and STs differ in cortical control (Paolone et al., 2013), plasma CORT response (Flagel et al., 2009), and dopamine response to Pavlovian cues, including within the PrL itself (Flagel et al., 2011; Pitchers et al., 2017), we expected to observe phenotypic differences in GR mRNA expression in the PrL. Yet, no significant differences were apparent. It should be noted, however, that GR mRNA expression may not be reflective of receptor (*i.e.*, protein) levels. Further investigation is warranted to determine if GR function in either the hippocampus or prelimbic cortex plays a role in incentive motivational processes. It is also possible that differences would have been apparent had GR mRNA been assessed in specific types of neurons (*e.g.*, tyrosine-hydroxylase positive or dopamine receptive) or within a given circuit (*e.g.*, PrL-nucleus accumbens). Ongoing studies are investigating this possibility. With these caveats in mind, the current data suggest that GRs, particularly within the ventral hippocampus, play a role in incentive motivational processes.

In conclusion, these findings establish that the neurobehavioral endophenotype associated with the propensity to sign-track does not include differences in stress-reactivity. Further, we provide additional evidence that glucocorticoids, which have primarily been implicated in aversive arousal (*but see* Deroche et al., 1993; Piazza et al., 1993), are also involved in appetitive motivation and, specifically, Pavlovian learning. In addition, expression of glucocorticoid receptors in the ventral hippocampus appear to be related to inherent cue-reward learning strategies. As these studies were limited to male rats, the role of corticosterone in cue-reward learning in females, and stress-reactivity in female goal-trackers and sign-trackers, warrants further investigation. Together, the findings reported here provide a foundation for future work to further examine the mechanism by which glucocorticoids interact with other neural systems to influence incentive motivational processes (*for further discussion see* Lopez & Flagel, 2020).

## Acknowledgments

We would like to thank Dr. Huda Akil and Dr. Stan Watson at the University of Michigan for providing the glucocorticoid receptor (GR) mRNA probe used for in situ hybridization and the gamma counter used to determine plasma corticosterone concentrations.

## Bibliography

Akil, H. (2005). Stressed and depressed. Nat Med, 11(2), 116–118. doi:10.1038/nm0205-116

Barr, J. L., Bray, B., & Forster, G. L. (2017). The Hippocampus as a Neural Link between Negative Affect and Vulnerability for Psychostimulant Relapse. In A. Stuchlik (Ed.), The Hippocampus: Plasticity and Functions (pp. 127–129). London, UK: BoD – Books on Demand, 2018.

Berridge, K. C., Robinson, T. E., & Aldridge, J. W. (2009). Dissecting components of reward: ‘liking’, ‘wanting’, and learning. Curr Opin Pharmacol, 9(1), 65–73. doi:10.1016/j.coph.2008.12.014

Butts, K. A., & Phillips, A. G. (2013). Glucocorticoid receptors in the prefrontal cortex regulate dopamine efflux to stress via descending glutamatergic feedback to the ventral tegmental area. International Journal of Neuropsychopharmacology, 16(8), 1799–1807. https://doi.org/10.1017/S1461145713000187

Campus, P., Covelo, I. R., Kim, Y., Parsegian, A., Kuhn, B. N., Lopez, S. A.,… Flagel, S. B. (2019). The paraventricular thalamus is a critical mediator of top-down control of cue-motivated behavior in rats. Elife, 8. doi:10.7554/eLife.49041

Clinton, S. M., Watson, S. J., & Akil, H. (2014). High novelty-seeking rats are resilient to negative physiological effects of the early life stress. Stress, 17(1), 97–107. https://doi.org/10.3109/10253890.2013.850670

Dallman, M. F., Akana, S. F., Strack, A. M., Scribner, K. S., Pecoraro, N., La Fleur, S. E.,… Gomez, F. (2004). Chronic stress-induced effects of corticosterone on brain: direct and indirect. Ann N Y Acad Sci, 1018, 141–150. doi:10.1196/annals.1296.017

Dallman, M. F., & Jones, M. T. (1973). Corticosteroid Feedback Control of ACTH Secretion: Effect of Stress-Induced Corticosterone Secretion on Subsequent Stress Responses in the Rat1. Endocrinology, 92(5), 1367–1375. https://doi.org/10.1210/endo-92-5-1367

Deroche-Gamonet, V., Sillaber, I., Aouizerate, B., Izawa, R., Jaber, M., Ghozland, S., Kellendonk, C., Le Moal, M., Spanagel, R., Schütz, G., Tronche, F., & Piazza, P. V. (2003). The Glucocorticoid Receptor as a Potential Target to Reduce Cocaine Abuse. The Journal of Neuroscience, 23(11), 4785–4790. https://doi.org/10.1523/JNEUROSCI.23-11-04785.2003

Deroche, V., Piazza, P. V., Deminiere, J. M., Le Moal, M., & Simon, H. (1993). Rats orally self-administer corticosterone. Brain Res, 622(1-2), 315–320. doi:10.1016/0006-8993(93)90837-d

Dorey, R., Pierard, C., Chauveau, F., David, V., & Beracochea, D. (2012). Stress-induced memory retrieval impairments: different time-course involvement of corticosterone and glucocorticoid receptors in dorsal and ventral hippocampus. Neuropsychopharmacology, 37(13), 2870–2880. doi:10.1038/npp.2012.170

Ehrman, R. N., Robbins, S. J., Childress, A. R., & O’Brien, C. P. (1992). Conditioned responses to cocaine-related stimuli in cocaine abuse patients. Psychopharmacology (Berl), 107(4), 523–529. doi:10.1007/bf02245266

Fanselow, M. S., & Dong, H. W. (2010). Are the dorsal and ventral hippocampus functionally distinct structures? Neuron, 65(1), 7–19. doi:10.1016/j.neuron.2009.11.031

Feldman, S., & Weidenfeld, J. (1993). The dorsal hippocampus modifies the negative feedback effect of glucocorticoids on the adrenocortical and median eminence CRF-41 responses to photic stimulation. Brain Res, 614(1-2), 227–232. doi:10.1016/0006-8993(93)91039-u

Feldman, S., & Weidenfeld, J. (1999). Glucocorticoid receptor antagonists in the hippocampus modify the negative feedback following neural stimuli. Brain Res, 821(1), 33–37. doi:10.1016/s0006-8993(99)01054-9

Fitzpatrick, C. J., Creeden, J. F., Perrine, S. A., & Morrow, J. D. (2016). Lesions of the Ventral Hippocampus Attenuate the Acquisition but Not Expression of Sign-Tracking Behavior in Rats. Hippocampus, 26(11), 1424–1434. doi:10.1002/hipo.22619

Fitzpatrick, C. J., Jagannathan, L., Lowenstein, E. D., Robinson, T. E., Becker, J. B., & Morrow, J. D. (2019). Single prolonged stress decreases sign-tracking and cue-induced reinstatement of cocaine-seeking. Behav Brain Res, 359, 799–806. doi:10.1016/j.bbr.2018.07.026

Flagel, S. B., Akil, H., & Robinson, T. E. (2009). Individual differences in the attribution of incentive salience to reward-related cues: Implications for addiction. Neuropharmacology, 56 Suppl 1, 139–148. doi:10.1016/j.neuropharm.2008.06.027

Flagel, S. B., Cameron, C. M., Pickup, K. N., Watson, S. J., Akil, H., & Robinson, T. E. (2011). A food predictive cue must be attributed with incentive salience for it to induce c-fos mRNA expression in cortico-striatal-thalamic brain regions. Neuroscience, 196, 80–96. doi:10.1016/j.neuroscience.2011.09.004

Flagel, S. B., Clark, J. J., Robinson, T. E., Mayo, L., Czuj, A., Willuhn, I.,… Akil, H. (2011). A selective role for dopamine in stimulus-reward learning. Nature, 469(7328), 53–57. doi:10.1038/nature09588

Flagel, S. B., & Robinson, T. E. (2017). Neurobiological Basis of Individual Variation in Stimulus-Reward Learning. Curr Opin Behav Sci, 13, 178–185. doi:10.1016/j.cobeha.2016.12.004

Flagel, S. B., Robinson, T. E., Clark, J. J., Clinton, S. M., Watson, S. J., Seeman, P.,… Akil, H. (2010). An animal model of genetic vulnerability to behavioral disinhibition and responsiveness to reward-related cues: implications for addiction. Neuropsychopharmacology, 35(2), 388–400. doi:10.1038/npp.2009.142

Garcia-Fuster, M. J., Flagel, S. B., Mahmood, S. T., Watson, S. J., & Akil, H. (2012). Cocaine withdrawal causes delayed dysregulation of stress genes in the hippocampus. PLoS One, 7(7), e42092. doi:10.1371/journal.pone.0042092

Harb, M. R., & Almeida, O. F. (2014). Pavlovian conditioning and cross-sensitization studies raise challenges to the hypothesis that overeating is an addictive behavior. Transl Psychiatry, 4, e387. doi:10.1038/tp.2014.28

Herman, J. P., Cullinan, W. E., Young, E. A., Akil, H., & Watson, S. J. (1992). Selective forebrain fiber tract lesions implicate ventral hippocampal structures in tonic regulation of paraventricular nucleus corticotropin-releasing hormone (CRH) and arginine vasopressin (AVP) mRNA expression. Brain Res, 592(1-2), 228–238. doi:10.1016/0006-8993(92)91680-d

Herman, J. P., Figueiredo, H., Mueller, N. K., Ulrich-Lai, Y., Ostrander, M. M., Choi, D. C., & Cullinan, W. E. (2003). Central mechanisms of stress integration: hierarchical circuitry controlling hypothalamo-pituitary-adrenocortical responsiveness. Front Neuroendocrinol, 24(3), 151–180. doi:10.1016/j.yfrne.2003.07.001

Herman, J. P., McKlveen, J. M., Ghosal, S., Kopp, B., Wulsin, A., Makinson, R.,… Myers, B. (2016). Regulation of the Hypothalamic-Pituitary-Adrenocortical Stress Response. Compr Physiol, 6(2), 603–621. doi:10.1002/cphy.c150015

Herman, J. P., McKlveen, J. M., Solomon, M. B., Carvalho-Netto, E., & Myers, B. (2012). Neural regulation of the stress response: glucocorticoid feedback mechanisms. Braz J Med Biol Res, 45(4), 292–298. doi:10.1590/s0100-879x2012007500041

Herman, J. P., Ostrander, M. M., Mueller, N. K., & Figueiredo, H. (2005). Limbic system mechanisms of stress regulation: hypothalamo-pituitary-adrenocortical axis. Prog Neuropsychopharmacol Biol Psychiatry, 29(8), 1201–1213. doi:10.1016/j.pnpbp.2005.08.006

Herman, J. P., Patel, P. D., Akil, H., & Watson, S. J. (1989). Localization and regulation of glucocorticoid and mineralocorticoid receptor messenger RNAs in the hippocampal formation of the rat. Mol Endocrinol, 3(11), 1886–1894. doi:10.1210/mend-3-11-1886

Hynes, T. J., Thomas, C. S., Zumbusch, A. S., Samson, A., Petriman, I., Mrdja, U.,… Lovic, V. (2018). Early life adversity potentiates expression of addiction-related traits. Prog Neuropsychopharmacol Biol Psychiatry, 87(Pt A), 56–67. doi:10.1016/j.pnpbp.2017.09.005

Kabbaj, M., Devine, D. P., Savage, V. R., & Akil, H. (2000). Neurobiological correlates of individual differences in novelty-seeking behavior in the rat: differential expression of stress-related molecules. J Neurosci, 20(18), 6983–6988.

Kawa, A. B., Bentzley, B. S., & Robinson, T. E. (2016). Less is more: prolonged intermittent access cocaine self-administration produces incentive-sensitization and addiction-like behavior. Psychopharmacology (Berl), 233(19-20), 3587–3602. doi:10.1007/s00213-016-4393-8

Kuhn, B. N., Campus, C., Flagel, S. B. (2018). Chapter 3: The neurobiological mechanisms underlying sign-tracking behavior. In T. Arthur, & J. Morrow (Ed.), Sign-tracking and drug addiction. Michigan Publishing, University of Michigan Library.

Liberzon, I., Lopez, J. F., Flagel, S. B., Vazquez, D. M., & Young, E. A. (1999). Differential regulation of hippocampal glucocorticoid receptors mRNA and fast feedback: relevance to post-traumatic stress disorder. J Neuroendocrinol, 11(1), 11–17.

Lister, R. G. (1987). The use of a plus-maze to measure anxiety in the mouse. Psychopharmacology (Berl), 92(2), 180–185. doi:10.1007/BF00177912

Lomanowska, A. M., Lovic, V., Rankine, M. J., Mooney, S. J., Robinson, T. E., & Kraemer, G. W. (2011). Inadequate early social experience increases the incentive salience of reward-related cues in adulthood. Behav Brain Res, 220(1), 91–99. doi:10.1016/j.bbr.2011.01.033

Lopez, S. A., & Flagel, S. B. (2020). A proposed role for glucocorticoids in mediating dopamine-dependent cue-reward learning. Stress, 1–41. doi:10.1080/10253890.2020.1768240

Lovic, V., Saunders, B. T., Yager, L. M., & Robinson, T. E. (2011). Rats prone to attribute incentive salience to reward cues are also prone to impulsive action. Behav Brain Res, 223(2), 255–261. doi:10.1016/j.bbr.2011.04.006

Maccari, S., Piazza, P. V., Deminiere, J. M., Angelucci, L., Simon, H., & Le Moal, M. (1991). Hippocampal type I and type II corticosteroid receptor affinities are reduced in rats predisposed to develop amphetamine self-administration. Brain Res, 548(1-2), 305–309.

Maggio, N., & Segal, M. (2007). Striking variations in corticosteroid modulation of long-term potentiation along the septotemporal axis of the hippocampus. J Neurosci, 27(21), 5757–5765. doi:10.1523/JNEUROSCI.0155-07.2007

Marchand, A. R., Barbelivien, A., Seillier, A., Herbeaux, K., Sarrieau, A., & Majchrzak, M. (2007). Contribution of corticosterone to cued versus contextual fear in rats. Behav Brain Res, 183(1), 101–110. doi:10.1016/j.bbr.2007.05.034

McKlveen, J. M., Myers, B., Flak, J. N., Bundzikova, J., Solomon, M. B., Seroogy, K. B., & Herman, J. P. (2013). Role of Prefrontal Cortex Glucocorticoid Receptors in Stress and Emotion. Biological Psychiatry, 74(9), 672–679. https://doi.org/10.1016/j.biopsych.2013.03.024

Meaney, M. J., Diorio, J., Francis, D., Widdowson, J., LaPlante, P., Caldji, C.,… Plotsky, P. M. (1996). Early environmental regulation of forebrain glucocorticoid receptor gene expression: implications for adrenocortical responses to stress. Dev Neurosci, 18(1-2), 49–72. doi:10.1159/000111395

Meyer, P. J., Lovic, V., Saunders, B. T., Yager, L. M., Flagel, S. B., Morrow, J. D., & Robinson, T. E. (2012). Quantifying individual variation in the propensity to attribute incentive salience to reward cues. PLoS One, 7(6), e38987. doi:10.1371/journal.pone.0038987

Mikics, E., Barsy, B., Barsvari, B., & Haller, J. (2005). Behavioral specificity of non-genomic glucocorticoid effects in rats: effects on risk assessment in the elevated plus-maze and the open-field. Horm Behav, 48(2), 152–162. doi:10.1016/j.yhbeh.2005.02.002

Morrow, J. D., Maren, S., & Robinson, T. E. (2011). Individual variation in the propensity to attribute incentive salience to an appetitive cue predicts the propensity to attribute motivational salience to an aversive cue. Behav Brain Res, 220(1), 238–243. doi:10.1016/j.bbr.2011.02.013

Morrow, J. D., Saunders, B. T., Maren, S., & Robinson, T. E. (2015). Sign-tracking to an appetitive cue predicts incubation of conditioned fear in rats. Behav Brain Res, 276, 59–66. doi:10.1016/j.bbr.2014.04.002

Paolone, G., Angelakos, C. C., Meyer, P. J., Robinson, T. E., & Sarter, M. (2013). Cholinergic control over attention in rats prone to attribute incentive salience to reward cues. J Neurosci, 33(19), 8321–8335. doi:10.1523/JNEUROSCI.0709-13.2013

Paxinos G WC (2007) The rat brain in stereotaxic coordinates. Burlington, MA, Academic Press.

Piazza, P. V., Deroche, V., Deminiere, J. M., Maccari, S., Le Moal, M., & Simon, H. (1993). Corticosterone in the range of stress-induced levels possesses reinforcing properties: implications for sensation-seeking behaviors. Proc Natl Acad Sci U S A, 90(24), 11738–11742. doi:10.1073/pnas.90.24.11738

Pitchers, K. K., Kane, L. F., Kim, Y., Robinson, T. E., & Sarter, M. (2017). ‘Hot’ vs. ‘cold’ behavioural-cognitive styles: motivational-dopaminergic vs. cognitive-cholinergic processing of a Pavlovian cocaine cue in sign- and goal-tracking rats. Eur J Neurosci, 46(11), 2768–2781. doi:10.1111/ejn.13741

Reul, J. M. H. M., Bosch, F. R. van den, & Kloet, E. R. de. (1987). Relative occupation of type-I and type-II corticosteroid receptors in rat brain following stress and dexamethasone treatment: Functional implications. Journal of Endocrinology, 115(3), 459–467. https://doi.org/10.1677/joe.0.1150459

Reul, J. M., & de Kloet, E. R. (1985). Two receptor systems for corticosterone in rat brain: microdistribution and differential occupation. Endocrinology, 117(6), 2505–2511. doi:10.1210/endo-117-6-2505

Rice, B. A., Eaton, S. E., Prendergast, M. A., & Akins, C. K. (2018). A glucocorticoid receptor antagonist reduces sign-tracking behavior in male Japanese quail. Exp Clin Psychopharmacol, 26(4), 329–334. doi:10.1037/pha0000195

Rice, B. A., Saunders, M. A., Jagielo-Miller, J. E., Prendergast, M. A., & Akins, C. K. (2019). Repeated subcutaneous administration of PT150 has dose-dependent effects on sign tracking in male Japanese quail. Exp Clin Psychopharmacol, 27(6), 515–521. doi:10.1037/pha0000275

Robinson, T. E., & Berridge, K. C. (1993). The neural basis of drug craving: an incentive-sensitization theory of addiction. Brain Res Brain Res Rev, 18(3), 247–291.

Robinson, T. E., & Flagel, S. B. (2009). Dissociating the predictive and incentive motivational properties of reward-related cues through the study of individual differences. Biol Psychiatry, 65(10), 869–873. doi:10.1016/j.biopsych.2008.09.006

Rodgers, R. J., Haller, J., Holmes, A., Halasz, J., Walton, T. J., & Brain, P. F. (1999). Corticosterone response to the plus-maze: high correlation with risk assessment in rats and mice. Physiol Behav, 68(1-2), 47–53.

Romeo, R. D., Ali, F. S., Karatsoreos, I. N., Bellani, R., Chhua, N., Vernov, M., & McEwen, B. S. (2008). Glucocorticoid receptor mRNA expression in the hippocampal formation of male rats before and after pubertal development in response to acute or repeated stress. Neuroendocrinology, 87(3), 160–167. doi:10.1159/000109710

Sandi, C., Loscertales, M., & Guaza, C. (1997). Experience-dependent facilitating effect of corticosterone on spatial memory formation in the water maze. Eur J Neurosci, 9(4), 637–642. doi:10.1111/j.1460-9568.1997.tb01412.x

Sarter, M., & Phillips, K. B. (2018). The Neuroscience of Cognitive-Motivational Styles: Sign- and Goal-Trackers as Animal Models. Behavioral Neuroscience, 132(1), 1–12. doi:10.1037/bne0000226

Saunders, B. T., & Robinson, T. E. (2010). A cocaine cue acts as an incentive stimulus in some but not others: implications for addiction. Biol Psychiatry, 67(8), 730–736. doi:10.1016/j.biopsych.2009.11.015

Saunders, B. T., & Robinson, T. E. (2011). Individual variation in the motivational properties of cocaine. Neuropsychopharmacology, 36(8), 1668–1676. doi:10.1038/npp.2011.48

Saunders, B. T., Yager, L. M., & Robinson, T. E. (2013). Cue-evoked cocaine “craving”: role of dopamine in the accumbens core. J Neurosci, 33(35), 13989–14000. doi:10.1523/JNEUROSCI.0450-13.2013

Shin, L. M., Orr, S. P., Carson, M. A., Rauch, S. L., Macklin, M. L., Lasko, N. B.,… Pitman, R. K. (2004). Regional cerebral blood flow in the amygdala and medial prefrontal cortex during traumatic imagery in male and female Vietnam veterans with PTSD. Arch Gen Psychiatry, 61(2), 168–176. doi:10.1001/archpsyc.61.2.168

Tomie, A., Aguado, A. S., Pohorecky, L. A., & Benjamin, D. (2000). Individual differences in pavlovian autoshaping of lever pressing in rats predict stress-induced corticosterone release and mesolimbic levels of monoamines. Pharmacol Biochem Behav, 65(3), 509–517.

Tomie, A., Silberman, Y., Williams, K., & Pohorecky, L. A. (2002). Pavlovian autoshaping procedures increase plasma corticosterone levels in rats. Pharmacol Biochem Behav, 72(3), 507–513.

Vanhille, N., Belin-Rauscent, A., Mar, A. C., Ducret, E., & Belin, D. (2015). High locomotor reactivity to novelty is associated with an increased propensity to choose saccharin over cocaine: new insights into the vulnerability to addiction. Neuropsychopharmacology, 40(3), 577–589. doi:10.1038/npp.2014.204

Walf, A. A., & Frye, C. A. (2007). The use of the elevated plus maze as an assay of anxiety-related behavior in rodents. Nat Protoc, 2(2), 322–328. doi:10.1038/nprot.2007.44

Walsh, R. N., & Cummins, R. A. (1976). The Open-Field Test: a critical review. Psychol Bull, 83(3), 482–504.

Yager, L. M., & Robinson, T. E. (2013). A classically conditioned cocaine cue acquires greater control over motivated behavior in rats prone to attribute incentive salience to a food cue. Psychopharmacology (Berl), 226(2), 217–228. doi:10.1007/s00213-012-2890-y

Zorawski, M., & Killcross, S. (2002). Posttraining glucocorticoid receptor agonist enhances memory in appetitive and aversive Pavlovian discrete-cue conditioning paradigms. Neurobiol Learn Mem, 78(2), 458–464. doi:10.1006/nlme.2002.4075

